# Structural streamlining of axonemal doublet microtubules in *Plasmodium* male gametes

**DOI:** 10.64898/2026.10.01.755548

**Authors:** Zhixun Li, Shuzhen Yang, Yujin Huang, Shanshan Ma, Liping Luo, Xiaoyang Hou, Kunyue Deng, Jianhua Wang, Guanbo Wang, Miao Gui, Jing Yuan, Qiang Guo

## Abstract

Male gametogenesis in malaria parasites produces up to eight flagellated gametes within approximately 15 min, requiring *de novo* construction of motile axonemes on an unusually short timescale. Here we combine cellular cryo-electron tomography, high-resolution cryo-EM, proteomics, and parasite genetics to define the molecular architecture and assembly states of the *Plasmodium* male gamete axoneme. The doublet microtubules retain the principal surface-associated motility modules but contain a markedly reduced luminal microtubule inner protein network. Their inner junction lacks PACRG and is instead organized by CFAP20 and CFAP52, whose combined loss disrupts doublet integrity and exflagellation. We further identify a calmodulin protein at the outer dynein arm docking region and show that outer dynein arms accumulate progressively during gametogenesis. These findings establish distinct form of axonemal remodeling in *Plasmodium* and reveal the ordered assembly of structural and motor modules during rapid male gamete formation.

## Main

Malaria remains a major global health burden caused by parasites of the genus *Plasmodium*^1^. Transmission depends on the sexual stages of the parasite, which are ingested by *Anopheles* mosquitoes during a blood meal and rapidly activated in the mosquito midgut^2^. Within approximately 15 min, each male gametocyte completes three rounds of genome replication and produces up to eight flagellated haploid male gametes^3^. Flagellar motility enables these short-lived gametes to encounter and fertilize female gametes and is therefore essential for transmission ^4^.

The male gamete is the only flagellated stage of the *Plasmodium* life cycle^5^. During gametogenesis, basal bodies form de novo at the nuclear envelope and nucleate axonemes in the cytoplasm before motile gametes emerge through exflagellation^3^. Unlike canonical basal bodies built from triplet microtubules, *Plasmodium* basal bodies contain singlet microtubules and undergo a singlet-to-doublet transition during axoneme formation^3,5^. Whereas ciliogenesis in many other eukaryotic cells typically unfolds over hours to days ^6–8^, *Plasmodium* constructs multiple motile axonemes within minutes. How a functional flagellum is built within this compressed developmental program remains unresolved.

Axonemal doublet microtubules (DMTs) provide the structural platform for motile cilia and flagella^9^. Dynein motors, regulatory complexes and radial spokes assemble on the DMT surface to drive and coordinate beating, whereas microtubule inner proteins (MIPs) pattern the lumen and reinforce the tubulin lattice and the junctions between the A- and B-tubules^10,11^. Structural comparisons have revealed substantial variation in this DMT-associated layer across cell types and evolutionary lineages ^12–14^. Thus, a conserved tubulin scaffold can accommodate distinct complements of structural, motor and regulatory proteins. So far, 18 of over 50 MIPs have been shown to be conserved across species whose MIP structures have been resolved, from unicellular species to mammals^12^. This raises the question of which minimal set of MIPs required to assemble a DMT in motile cilia and flagella.

Specialized flagella illustrate how axonemal architecture can be tuned in different directions ^15^. Eel sperm flagella, which contain a 9+0 axoneme and lack the central pair and several major canonical motility-associated complexes, have been described as a molecular minimum system for motile cilia, although the MIP repertoire has not been revealed due to limited resolution ^16^. By contrast, trypanosomatid parasites retain 9+2 axonemes but show lineage-specific elaboration of DMT-associated proteins^17,18^. These examples suggest that axonemes can be minimized, elaborated, or remodeled according to cellular demand. *Plasmodium* presents a different biological problem: its axonemes are assembled rapidly, without intraflagellar transport, and function only during the brief male-gamete stage^3,5^. The molecular composition of its DMTs, the structural adaptations that preserve DMT integrity and the sequence in which major axonemal modules appear during gametogenesis are unknown.

Here, by leveraging cryo-electron tomography with high-resolution cryo-EM, mass spectrometry, and reverse genetics, we resolve the axonemal architecture of intact male gametes and activated gametocytes. We show that *Plasmodium* retains the principal surface-associated motility machinery but has a reduced luminal MIP network and a PACRG-independent inner junction. CFAP20 and CFAP52 act cooperatively to maintain DMT integrity, while orthogonal structural, proteomic and localization analyses identify a calmodulin protein at the outer dynein arm docking region. Time-resolved cryo-ET further reveals early inner-junction formation followed by progressive ODA assembly. Together, these findings define a reduced but specialized DMT architecture associated with rapid axoneme assembly and male gamete motility.

## Results

### Native architecture of the *Plasmodium* male gamete axoneme

To determine the native organization of the *Plasmodium* male gamete axoneme, we induced exflagellation of the *Plasmodium yoelii* (*P. yoelii*) male gametocytes (Fig. 1a,b) and vitrified intact male gametes for cryo-electron tomography (cryo-ET). Tomographic reconstructions revealed axonemes containing a central pair of singlet microtubules surrounded by nine DMTs (Fig. 1c-d). Subtomogram averaging further resolved prominent protein assemblies on the DMT surface, whereas the DMT lumen contained markedly fewer internal densities than previously described motile cilia and flagella^10,14,18–20^ (Fig. 1e, Fig. S1 and S2). The central pair contained a bridge connecting the C1 and C2 microtubules, together with at least 11 morphologically distinct central-pair projections with 16-nm or 32-nm periodicities (Fig. 1f). Thus, the rapidly assembled male gamete flagellum contains an ordered central apparatus ^21,22^ and extensive surface-associated machinery despite its sparse luminal protein network.

**Fig. 1.**
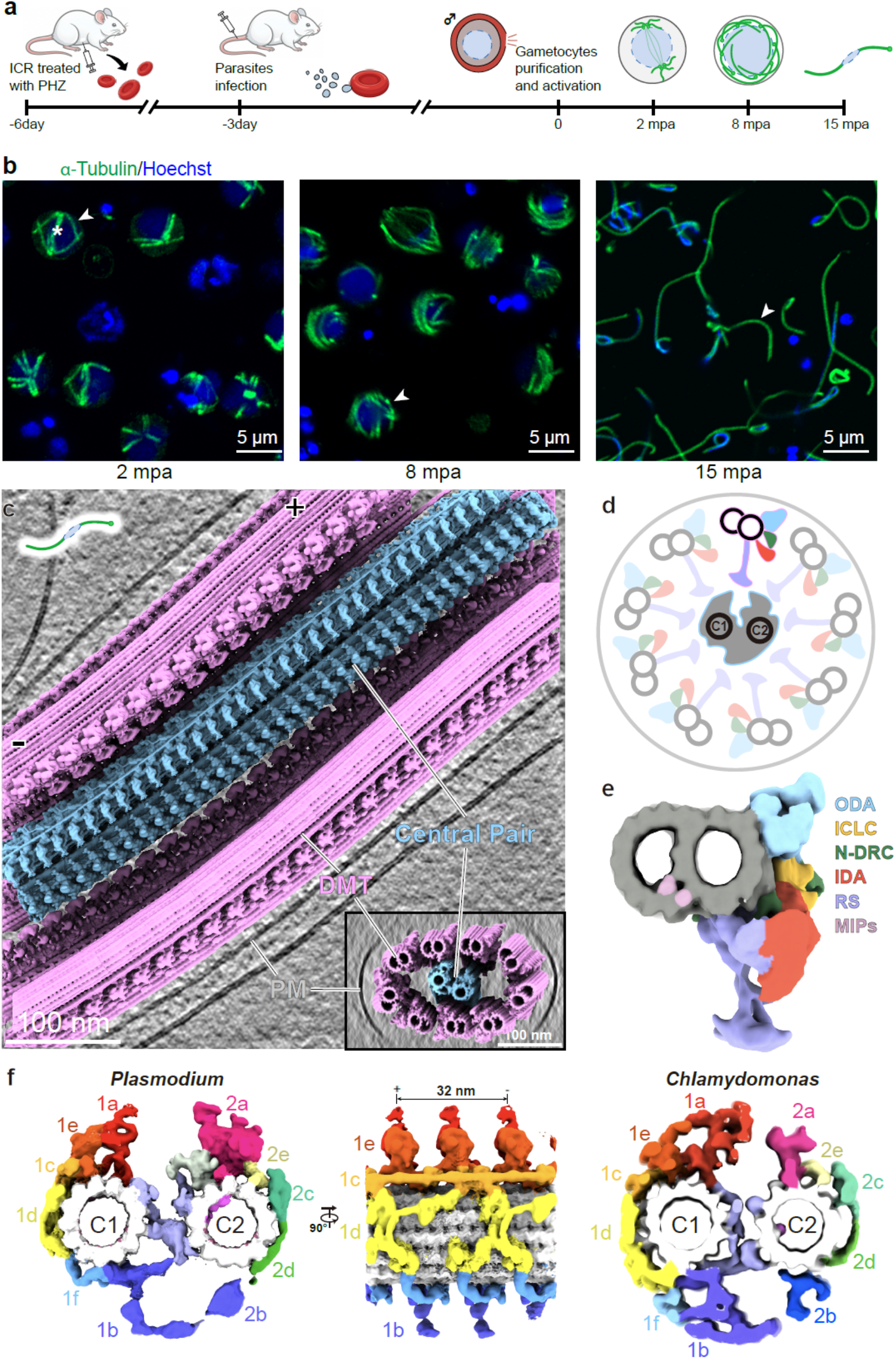
In situ architecture of the *Plasmodium* male gamete axoneme. **a.** Experimental workflow for inducing *Plasmodium* gametocytes in mice, followed by purification of male gametocytes and male gametes for imaging. Red, erythrocytes; gray, asexual-stage parasites invading erythrocytes; blue, nuclei. Green in the cytoplasm, axonemes; green within the nuclei, spindle microtubules. **b.** Immunofluorescence analysis of cytoplasmic axoneme formation in male gametocytes at 2, 8 and 15 min post-activation (mpa). Parasites were stained with anti-α-tubulin antibody and Hoechst 33342. Arrowheads indicate axonemes; the asterisk indicates spindle microtubules. **c.** Three-dimensional rendering of a representative cryo-electron tomogram showing the axoneme of a male gamete. The plasma membrane (PM), doublet microtubules (DMTs) and central pair are annotated. Two DMTs were omitted from the rendering to reveal the central pair. Inset displays the cross-section view of the axoneme. **d.** Schematic cross-section of the axoneme, showing the central pair and nine surrounding DMTs. **e.** Cross-section view of *Plasmodium* DMT structures resolved by subtomogram averaging. **f.** Cross-section and longitudinal views of *Plasmodium* central pair complex resolved by subtomogram averaging. The 32-nm averaged structure reveals C1, C2 microtubules (white) and associated densities (1a-f, 2a-e). The central pair of *Chlamydomonas reinhardtii* flagella (EMD-31143) is shown for comparison and colored accordingly.

### *Plasmodium* DMTs retain core motility modules while reducing luminal MIPs

To define the organization of the motility machinery, we resolved the 96-nm repeat of the *Plasmodium* DMT (Fig. 2a, Table S1). The principal force-generating and regulatory assemblies were retained ^23–26^: outer and inner dynein arms (ODAs and IDAs) decorated the A-tubule to drive microtubule sliding, the nexin-dynein regulatory complex (N-DRC) linked neighboring DMTs to constrain sliding into bending, and RSs extended toward the central pair to couple central-pair signals to dynein regulation (Fig. 2a). We also observed an intermediate chain/light-chain (ICLC) complex associated with the ODA and A-tubule (Fig. 2a). In the current map, this ICLC density does not contact the IDA in *Plasmodium*, in contrast to the outer-inner dynein linker described in some other axonemes^27,28^.

**Fig. 2.**
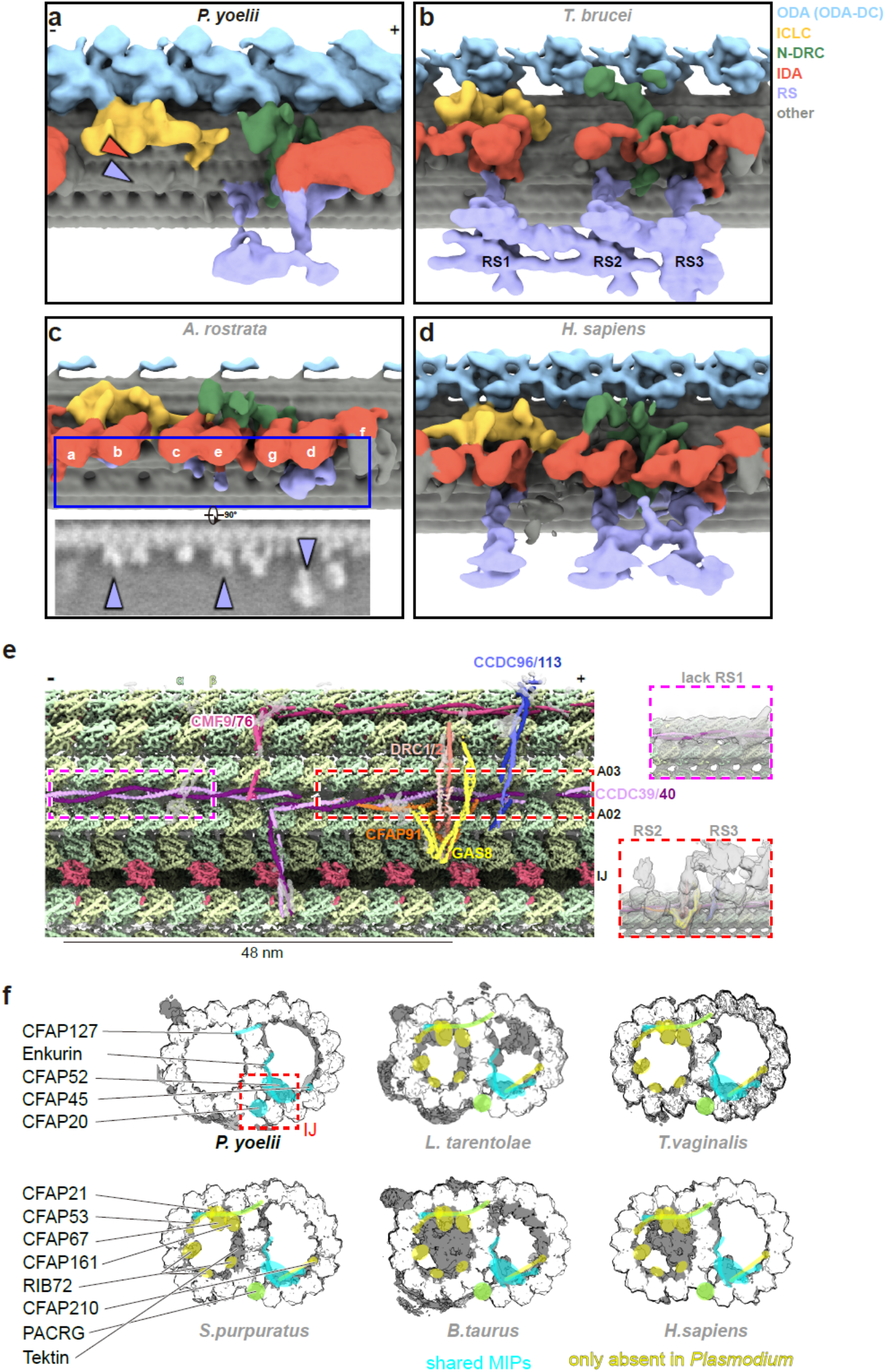
Conservation and reduction of DMT-associated assemblies in *Plasmodium*. **a-d**. Comparison of 96-nm repeat architecture of DMT-associated motor and regulatory assemblies in *Plasmodium yoelii* (subtomogram averaged map), *Trypanosoma brucei* (EMD-47524), *Anguilla rostrata* (EMD-45870) and *Homo sapiens* (EMD-27462). ODA, fIC/LC, N- DRC, IDA, RS are color-coded. Different IDA subunits (a, b, c, e, g, d, f) are labeled in (c). Orange and purple arrowheads indicate regions corresponding to reduced or absent IDA-associated densities in *Plasmodium*, respectively (a). The inset in (c) shows a 1-nm-thick longitudinal slice of eel DMT and purple arrows highlight the residual RS density. (ODA: outer dynein arm; ICLC: Intermediate chain/light chain; N-DRC: nexin-dynein regulatory complex; IDA: inner dynein arm; RS: radial spoke). **e**. View of the *Plasmodium* DMT (cryo-EM map) from the radial spoke side, with conserved 96-nm repeat assemblies color-coded and labeled. The regions corresponding to the RS1 base are boxed in purple and displayed with the density map of RS in right inset, whereas the regions corresponding to the RS2/3 base are boxed in red and displayed similarly in right inset. **f**. Comparison of 48-nm repeat MIPs organization across resolved DMT structures from *Leishmania tarentolae*, EMD-47661; *Trichomonas vaginalis*, EMD-46580; *Strongylocentrotus purpuratus*, EMD-40619; *Bos taurus*, EMD-17187; *Homo sapiens*, EMD-35810. IJ, inner junction. Blue: MIPs conserved across the compared species. Yellow: MIPs present in other species but absent from *Plasmodium*.

We next compared the 96-nm repeat architecture of *Plasmodium* DMTs with those from other specialized motile systems, including the trypanosomatid parasite (*Trypanosoma brucei*)^17^, eel (*Anguilla rostrata*) sperm^16^ and human (*Homo sapiens*) sperm flagella^29^ (Fig. 2b-d). These examples illustrate distinct modes of axonemal specialization.

Despite this overall conservation, the 96-nm repeat was selectively reduced. RS2 and RS3 were present, whereas RS1 and adjacent IDA-associated densities were absent or strongly reduced (Fig. 2a). Because CCDC39/40 helps establish the 96-nm periodicity of radial spokes and inner dynein arms ^30,31^, we examined whether this scaffold was retained. Coiled-coil densities consistent with CCDC39/40 were present at the radial-spoke base (Fig. 2e), indicating that the 96-nm repeat is maintained despite the absence of RS1 and reduction of nearby dynein-associated densities. Together with comparative genomic analyses showing that *Plasmodium* encodes a reduced repertoire of IDA- and RS1-associated components^32,33^, these data suggest selective simplification of specific motility modules rather than loss of the underlying periodic scaffold.

The contrast between extensive surface decoration and a sparse luminal protein network prompted us to determine which structural and motility-associated modules are retained, reduced or remodeled in the *Plasmodium* axoneme. We next determined high-resolution structures of *Plasmodium* DMTs (Fig. 2f, Fig. S3, Table S1). To preserve native axonemal organization, exflagellated male gametes were plunge-frozen without biochemical purification, and DMT particles were picked directly from the crowded cellular environment using a semi-automated workflow supported by GisSPA^34^. The central ribbon region of the DMT was resolved to 3.28 Å resolution (Fig. S3), revealing markedly fewer luminal densities than in previously resolved motile cilia^18–20,35^ (Fig. 2f).

### A PACRG-independent inner junction stabilizes *Plasmodium* DMTs

To define the molecular basis of this reduced scaffold, we reconstructed the complete 48-nm DMT repeat by combining the 16-nm periodicity of the inner-junction region with the 24-nm periodicity of the ODA region through sequential focused classification (Fig. S3). Because many *Plasmodium* axonemal proteins are sequence-divergent, assignments were based on convergent evidence from side-chain-resolved density, AlphaFold-predicted models and proteins detected by mass spectrometry in male-gamete-enriched samples (Fig. S4 and Fig. S5). The resulting map lacked tektin-like longitudinal bundles and densities corresponding to NME7 (CFAP67), CFAP161 and RIB72 (EFHC1/EFHC2) (Fig. 2f, Fig. 3a). Sequence searches and male-gamete proteomic datasets likewise identified no clear *P. yoelii* orthologs of these proteins, supporting a genuine reduction of the MIP repertoire rather than loss of density through limited resolution. The A-tubule lumen contained few globular MIP densities, and instead, the remaining internal decoration was dominated by long α-helical densities positioned along inter-protofilament clefts. At a region occupied by CFAP53, CFAP141 and MNS1 in other axonemes^36^, *Plasmodium* lacked identifiable CFAP53 and CFAP141 orthologs but contained MNS1-related density. MNS1 occupied multiple sites with a 16-nm offset and extended across the A- and B-tubule interface (Fig. 3a and Fig. S6a). Thus, a retained helical MIP is redeployed across positions occupied by several proteins in other species, reinforcing selected regions of the reduced DMT scaffold.

**Fig. 3.**
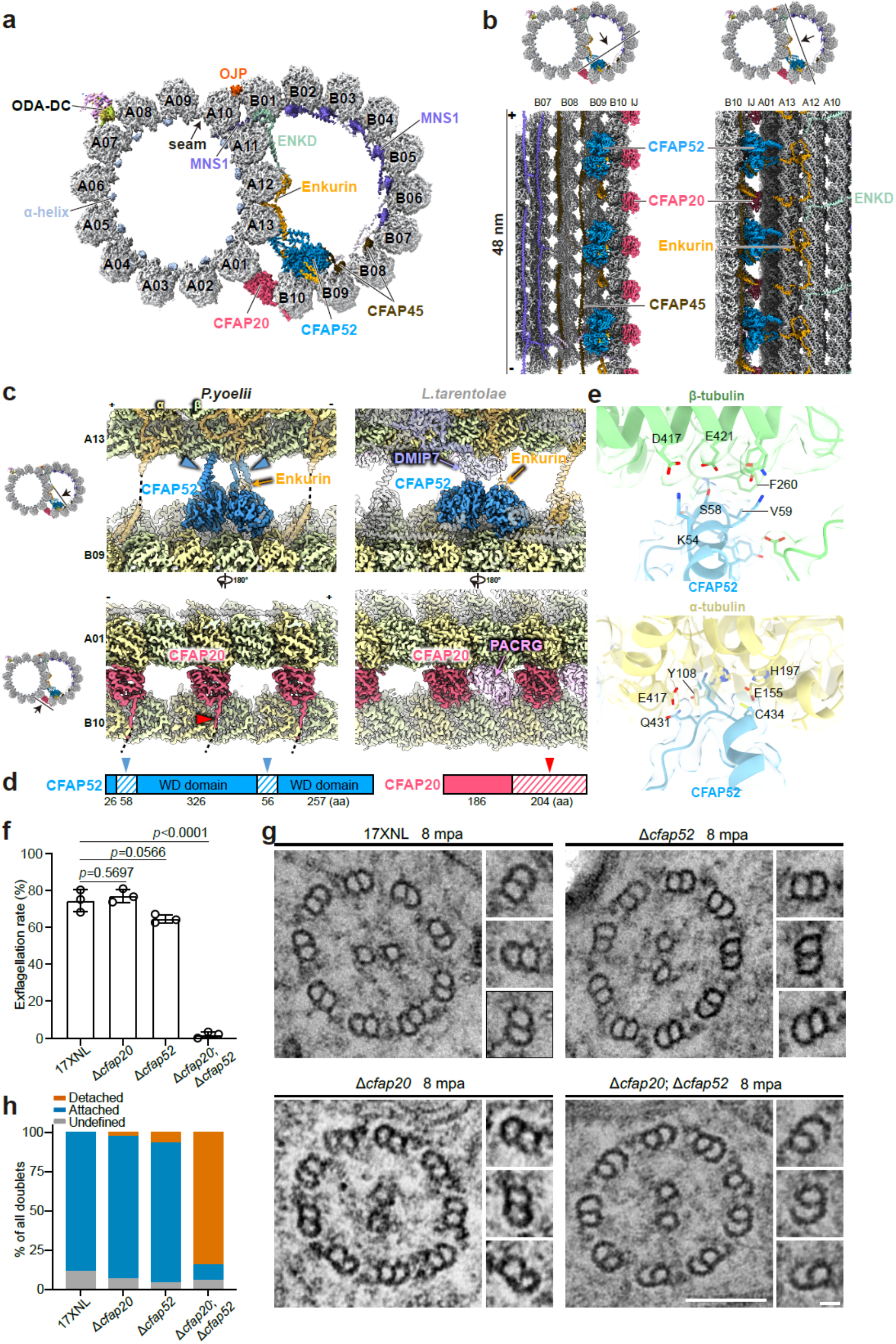
48-nm repeat of DMT reveals a PACRG-independent inner junction in *Plasmodium*. **a.** Cryo-EM map shows the cross-section view of *Plasmodium* DMT, with protofilaments numbered and MIPs annotated. **b.** Longitudinal sections of the DMT structure, all MIPs are color-coded and labeled. Corresponding cross sections are shown with viewing positions and angles indicated by black lines and arrows. **c.** Structural comparison of CFAP52 and CFAP20 at the inner junction of *Plasmodium.yoelii* and *Leishmania.tarentolae* (EMD-47661). The additional densities unique to *Plasmodium* are indicated by red arrowheads. **d.** Sequence insertions corresponding to the additional densities in Plasmodium CFAP20 and CFAP52. Hatched regions indicate insertions absent from the compared orthologs; red arrowheads indicate regions corresponding to additional densities in the cryo-EM map. **e.** Interface between the CFAP52 additional regions and α-/β-tubulins. Side chains involved in the interface are shown and annotated. **f.** In vitro exflagellation assay of male gametocytes from the indicated parasite lines. Data are mean ± SD from the infected blood of three mice in each group, statistical analysis was performed using two-tailed t-test. A representative result from three independent experiments is shown. **g.** TEM of axonemal microtubules architecture in the gametocytes at 8 mpa. Scale bars, 100 nm. Insets show enlargements of the doublet microtubules from the corresponding images. Scale bars, 20 nm. Images are representative of three independent experiments. **h.** Frequency of DMT structural states quantified from TEM sections.

The B-tubule also contained a limited set of long α-helical MIPs. MNS1 and CFAP45 bound adjacent inter-protofilament clefts and established a 48-nm periodicity along the B-tubule (Fig. 3a,b and Fig. S6b). At the outer junction, we observed two densities, here assigned to Enkurin domain-containing protein (ENKD) across A12-A11-B01 and outer junction protein (OJP) across A10-B01, that connected the A- and B-tubules (Fig. 3a). Across several previously resolved DMT structures, the equivalent outer-junction position across A12-A11-B01 is occupied by CFAP77 ^37,38^. In *Plasmodium*, however, no clear CFAP77 orthologue was identified, and the ENKD density, although similarly positioned, differed in shape from CFAP77 (Fig. S6c). These observations suggest that the outer junction is also remodeled in *Plasmodium* DMTs, potentially through divergent components rather than by the canonical CFAP77 architecture.

The most distinctive feature was found at the inner junction. In previously characterized axonemes, CFAP20 and PACRG alternate along the inner junction and together support B-tubule closure and ciliary motility^10,19,39^. By contrast, *Plasmodium* DMTs contained CFAP20, CFAP52 and Enkurin at the inner junction but lacked PACRG (Fig. 3b,c). Among structurally characterized motile axonemes, *Plasmodium* therefore provides an example of a naturally PACRG-independent inner junction. This finding is notable because PACRG loss disrupts B-tubule integrity and impairs motility in other organisms ^18,40^, implying that *Plasmodium* stabilizes the inner junction through an alternative architecture.

Consistent with this idea, *Plasmodium* CFAP20 and CFAP52 contained parasite-specific features at their tubulin interfaces. Additional density associated with the CFAP52 WD-repeat domain contacted the A-tubule, whereas a C-terminal extension of CFAP20 contacted the B-tubule (Fig. 3c-d). Structural analysis indicated that the contacts between tubulins and the additional CFAP52 region are mainly mediated by hydrogen bonds and hydrophobic interactions (Fig. 3e). Sequence alignments mapped these densities to insertions in *Plasmodium* CFAP20 and CFAP52 (Fig. S7 and S8). The extensions enlarge the interfaces of both proteins with the DMT lattice and reinforce the PACRG-independent inner junction.

To test the functional contribution of CFAP20 and CFAP52, we generated three *P. yoelii* mutant lines in which CFAP20, CFAP52, or both proteins were no longer expressed. Loss of either protein alone caused no detectable change in exflagellation or DMT architecture under the conditions examined (Fig. 3f-h). This contrasts with observations in trypanosomatids, where disruption of inner-junction components, including CFAP20 or CFAP52, impairs flagellar motility^18^. Simultaneous loss of both CFAP20 and CFAP52 proteins, however, disrupted DMT integrity and impaired male gamete exflagellation (Fig. 3f). Transmission electron microscopy at 8 mpa further revealed disrupted, detached or incompletely closed B-tubules in the double mutant, whereas DMT profiles in either single mutant remained comparable to those in wild-type parasites (Fig. 3g,h). The absence of a strong single-mutant phenotype and the marked double-mutant defect demonstrate that CFAP20 and CFAP52 make cooperative, partially redundant contributions to DMT integrity. Together, the structural and genetic phenotypes establish a PACRG-independent inner junction architecture.

### A specific calmodulin protein associates with the *Plasmodium* ODA docking region

The ODAs provide a major source of force for axonemal beating and are anchored to the A-tubule by the ODA docking complex (ODA-DC)^41^. In *Plasmodium* DMT map, the ODA docking region contained DC1-, DC2- and DC3-like densities (Fig. 4a,b and Fig. S4b), broadly resembling the tripartite docking complex described in *Chlamydomonas reinhardtii*^37,42^. In addition, the *Plasmodium* ODA-DC contained an additional ring-shaped density surrounding DC1- and DC2-like densities, indicating that the docking region is structurally modified in this parasite.

**Fig. 4.**
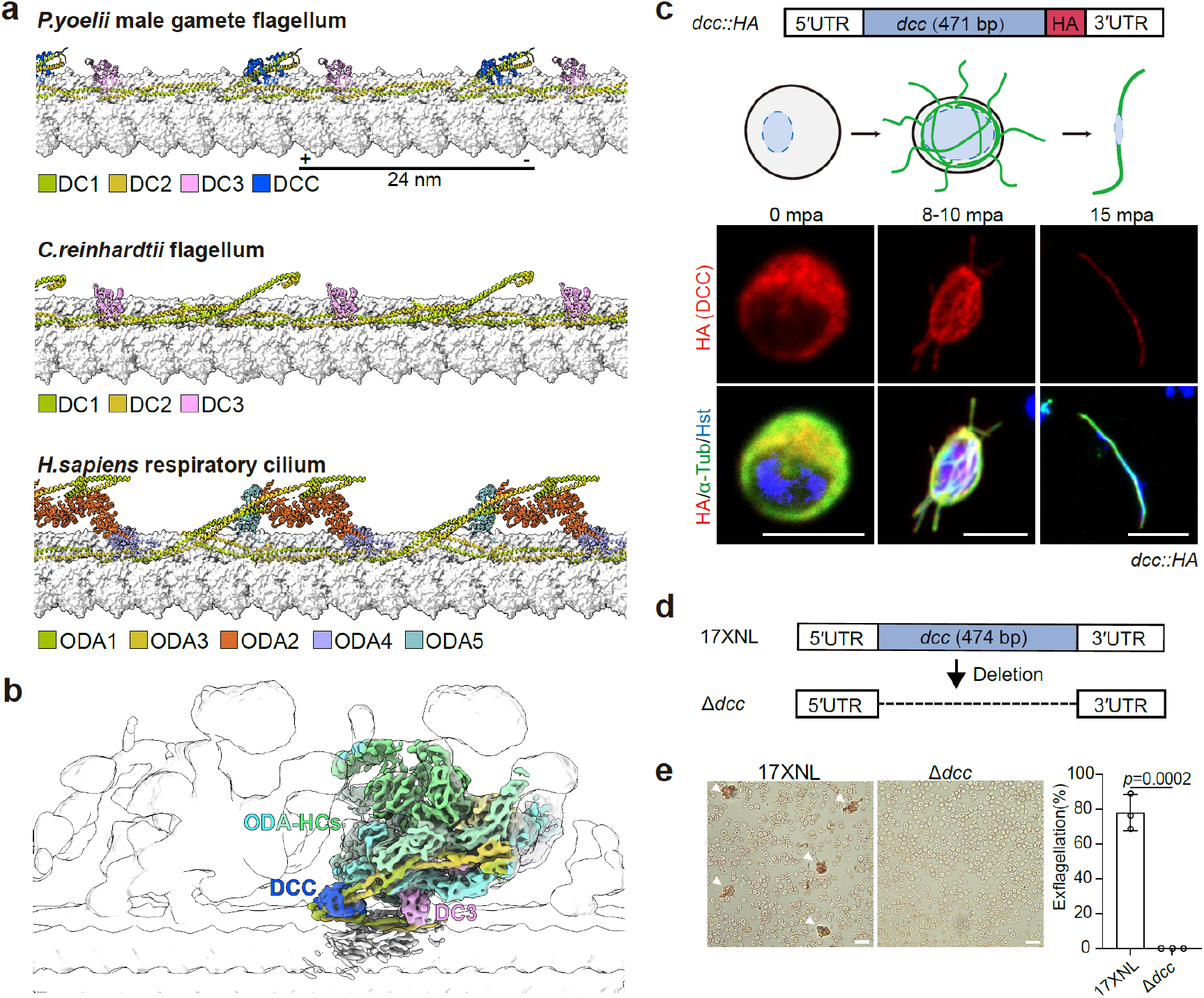
A specific calcium-binding protein associates with the *Plasmodium* ODA docking complex. **a.** Atomic model comparison of the *Plasmodium* ODA docking complex (ODA-DC) with the ODA-DCs from *Chlamydomonas reinhardtii* (PDB: 6U42) and *Homo sapiens* respiratory cilium (PDB: 7UNG). **b.** Cryo-EM of the calcium-binding protein (DCC) in complex with the surrounding ODA heavy chains (ODA-HCs). Three ODA-HCs are colored individually. The transparent density corresponds to the consensus map, while the colored density corresponds to the focused-refined composite map. **c.** Immunofluorescence analysis of HA tagged DCC and α-Tubulin expression during the male gametogenesis of the *dcc::HA* parasite line. Upper panel: the CRISPR-Cas9 strategy used to modify the *dcc* locus. Middle panels: schematic representation of male gametogenesis development with the axoneme in green and the nucleus in blue. Images are representative of three independent experiments. Scale bars, 5 μm. **d.** Schematic showing the *dcc* gene deletion in the *P. yoelii* 17XNL parasite. The 474 bp coding sequence of dcc was deleted to generate mutant clones. **e.** In vitro exflagellation assay of male gametocytes. White arrowheads indicate the exflagellation centers of activated male gametocytes. Representative images from three independent experiments are shown. Scale bars, 20 μm. Quantification shows male gametocyte exflagellation, data are mean ± SD from the infected blood of three mice in each group, statistical analysis was performed using a two-tailed t-test.

To identify this additional density, we analyzed male gamete-enriched fractions by mass spectrometry and searched for candidate proteins compatible with the density shape and predicted structure (Fig. S4, S9a). Additionally, *de novo* model building using CryoAtom^43^, followed by sequence search, was performed (Fig. S9b). These analyses assigned the ring-shaped density to a calmodulin protein encoded by UniProt ID A0A078K5I3. Endogenous HA tagging confirmed that this protein localizes to axonemes during gametogenesis (Fig. 4c). Because *Plasmodium* encodes several calmodulin or calmodulin-like proteins, we designate A0A078K5I3 docking-complex calmodulin (DCC).

The structural position of the DCC suggests an accessory role in ODA regulation or stabilization rather than a primary role in ODA anchoring. DCC encircled DC1 and DC2 and contacted the ODA heavy chain only locally, whereas the DC3-like density and coiled-coil docking scaffold formed more extensive interfaces with the A-tubule and ODA (Fig. 4b).

Deletion of *dcc* impaired male gamete exflagellation (Fig. 4d,e and Fig. S9c-h), supporting a functional contribution to flagellum formation or motility. Together, the structural assignment, axonemal localization and mutant phenotype identify DCC as a parasite-specific, functionally important component of the ODA docking complex.

### Time-resolved cryo-ET reveals progressive ODA assembly during male gametogenesis

We next used time-resolved cryo-ET to determine when the inner junction and ODAs assemble during male gametogenesis. Based on our previous work^3^, activated male gametocytes form two tetrads of basal bodies by 2 min post activation (mpa), from which axonemal DMTs begin to elongate. We vitrified independently activated samples at 2 and 8 mpa and compared them with mature male gametes collected at 15 mpa (Fig. 1a,b), capturing successive states of axoneme construction.

At 2 mpa, many DMTs had not yet organized around a mature central-pair scaffold (Fig. 5a, Fig. S10). This indicates that early axoneme initiation in *Plasmodium* can proceed before complete organization of the mature axonemal geometry. Subtomogram classification showed that most early DMT segments already contained inner-junction densities corresponding to CFAP20 and CFAP52 (Fig. S11a,b). The early incorporation of these components establishes the A-B tubule interface during DMT formation.

**Fig. 5.**
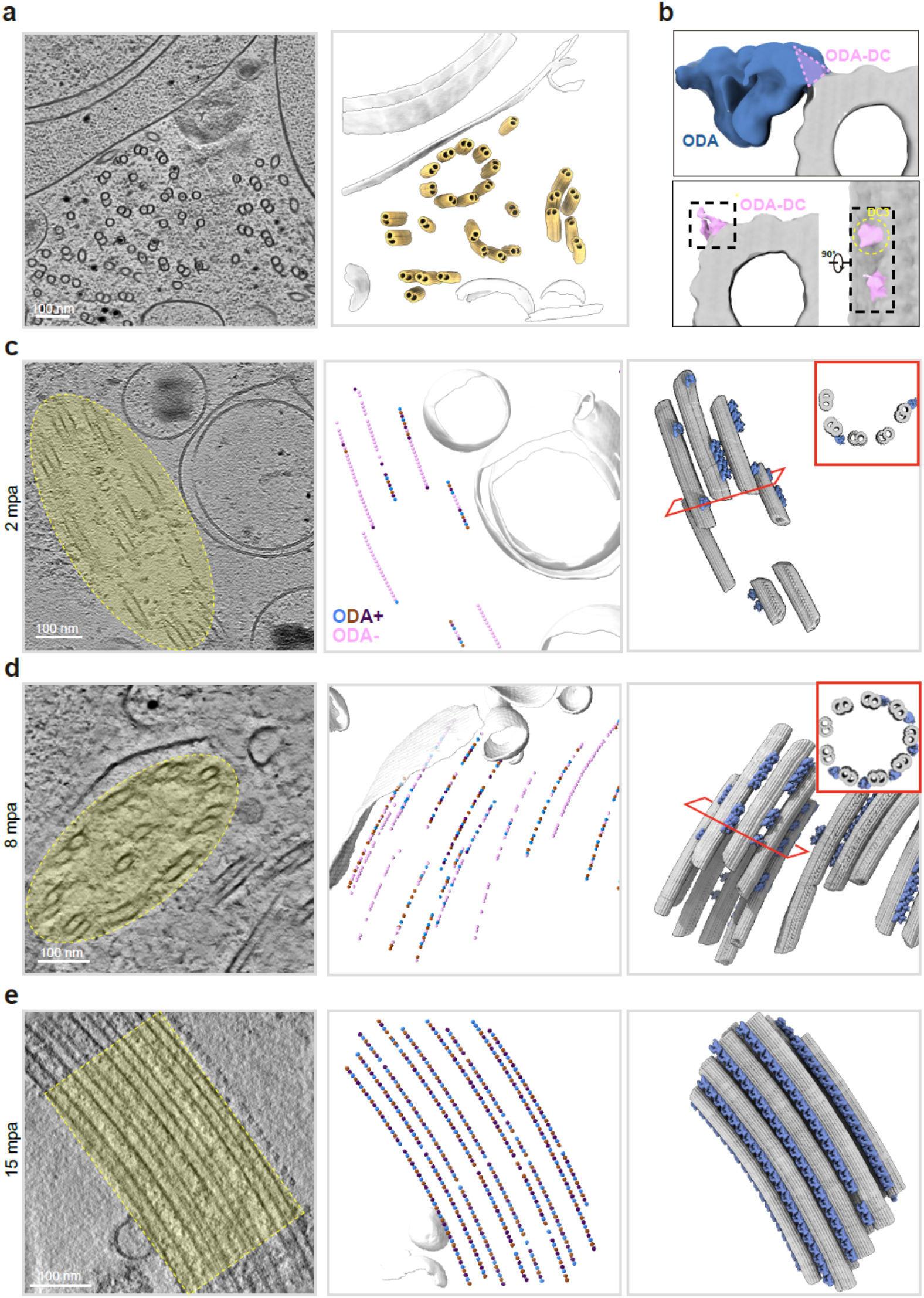
Time-resolved cryo-ET reveals progressive ODA assembly on DMTs. **a.** Three-dimensional rendering of a *P. yoelii* male gametocyte activated for 2 minutes. Doublet microtubules (DMTs) are colored yellow. **b**. Subtomogram average of DMTs with assembled ODA (ODA+, left) and before ODA assembled (ODA-, right). The dashed boxes highlight the ODA-DC region in cross-sectional and side views. Density of DC3 indicated in yellow circle. **c-e.** Representative tomograms showing ODA assembly at different time points after activation (c, 2 mpa; d, 8mpa; e, 15 mpa). **Left:** Tomogram slices (thickness: 1.4 nm) showing axonemes arrangement. The yellow regions indicate microtubules assembled into axonemal arrangement. Scale bar, 100 nm. **Middle:** Three-dimensional spatial distribution of DMTs with assembled ODA (ODA+) and without ODA (ODA-). ODA+ are colored blue, brown and purple according to 3D classification results, whereas ODA-doublets are colored pink. **Right:** Three-dimensional rendering shows the fine structure and spatial distribution of ODA on DMTs. The inset displays the cross-section view of DMTs before axoneme formed 9+2 arrangement.

By contrast, ODA occupancy was heterogeneous at 2 mpa. Only approximately 39.9% of DMT regions contained clear ODA density, whereas many regions contained the ODA docking complex without a fully assembled ODA (Fig. 5b). ODA-positive segments occurred in discrete clusters at multiple positions along individual DMTs, consistent with cooperative assembly of neighboring motors into axonemal arrays. From 2 to 8 mpa, the proportion of ODA-decorated DMT regions increased progressively (Fig. 5c-e), indicating continued motor incorporation during male gametogenesis.

Together, the time-resolved structures reveal a modular assembly sequence. Inner-junction components are incorporated early to establish the A-B tubule interface, whereas ODA occupancy increases progressively as gametogenesis proceeds. *Plasmodium* therefore constructs the DMT scaffold before completing its decoration with force-generating motors, providing a structural mechanism for coordinating axoneme assembly within the short period of male gamete formation.

## Discussion

Our structural, proteomic, localization and genetic analyses define an axonemal architecture that differs from both canonical motile cilia and other specialized parasite flagella. *Plasmodium* retains an ordered central apparatus and the principal DMT-surface machinery for force production and regulation, while sharply reducing the luminal MIP network (Movie S1). This asymmetric reduction separates two layers of axonemal organization: a broadly retained motility apparatus and a highly adaptable internal scaffold.

Comparisons of previously determined DMT structures identify a core set of at least 18 recurrent MIPs, despite extensive variation among cell types and species^12^. *Plasmodium* lowers this apparent conserved minimum substantially. Its DMTs lack several recurrent A-tubule MIPs and tektin-like bundles, while retaining or redeploying a small subset of helical MIPs and junctional contacts. This reduction differs from that in eel sperm, which lack the central pair and several major canonical motility-associated complexes ^16^, and from the lineage-specific elaboration of trypanosomatid DMTs ^17,18^. *Plasmodium* therefore represents a distinct solution: it preserves the major external motility apparatus while reducing and reorganizing the DMT luminal scaffold. The retained MIPs are concentrated at lattice clefts and A–B-tubule interfaces, suggesting that the reduced scaffold preserves the interactions most critical for DMT closure and stability. This architecture provides a plausible structural solution for constructing multiple functional axonemes within minutes: reducing luminal complexity while maintaining the junctional contacts required for rapid assembly.

The PACRG-independent inner junction illustrates how *Plasmodium* compensates for reduction rather than merely tolerating it. In other organisms, CFAP52, PACRG and CFAP20 alternate along the inner junction and are important for B-tubule closure and motility^10,40,44,45^. In *Plasmodium*, expanded CFAP20- and CFAP52-tubulin interfaces replace this arrangement, and the double-mutant phenotype demonstrates that these proteins form a cooperative stabilizing system. The lack of strong phenotypes in either single mutant further indicates that robustness is distributed across the remodeled interface. Thus, streamlining is accompanied by reinforcement of the contacts that remain.

Time-resolved cryo-ET further reveals the order in which major DMT modules assemble. Inner-junction densities are prevalent in early DMT segments, whereas ODA occupancy increases between early gametocytes and mature male gametes. This sequence establishes the A-B tubule interface before complete motor decoration and shows that *Plasmodium* axoneme formation proceeds through coordinated structural and motor-assembly phases. Such an assembly strategy may help *Plasmodium* balance speed and function: the parasite first establishes a stable DMT scaffold and then progressively decorates it with force-generating machinery.

The identification of a specific calmodulin protein DCC at the ODA docking region adds another *Plasmodium*-specific feature to this architecture. DCC surrounds DC1 and DC2 but makes only limited contact with the ODA heavy chain, suggesting a regulatory or stabilizing role rather than a primary anchoring role. The exflagellation defect in the *dcc* mutant supports a contribution to male gamete flagellum function, but the precise mechanism remains to be determined. Given the importance of calcium signaling in gametocyte activation and flagellar motility^46–48^, DCC may provide a route through which calcium-binding proteins are integrated into the ODA docking machinery.

Together, our findings establish *Plasmodium* male gametes as a distinct example of axonemal remodeling: a reduced luminal scaffold is combined with reinforced junctional contacts, a parasite-specific ODA-docking component and staged incorporation of structural and motor modules. The work links molecular structures to proteomic identification, cellular localization and genetic phenotypes, and thereby moves beyond a descriptive inventory of the parasite axoneme. More broadly, it shows how the DMT-associated protein layer can vary while preserving a functional motility apparatus during an exceptionally compressed developmental program. As male-gamete motility is indispensable for fertilization and mosquito transmission, the divergent structural features identified across the *Plasmodium* axoneme provide a molecular framework for developing transmission-blocking strategies that selectively disrupt flagellum assembly, stability or motility.

## Methods

### Mice and mosquitoes usage and ethics statement

The animal experiments conducted in this study were approved by the Committee for the Care and Use of Laboratory Animals of Xiamen University (approval number: XMULAC20190001). Female ICR mice, aged 5–6 weeks, were obtained from the Animal Care Center of Xiamen University. The mice were housed under controlled environmental conditions at 22–24 °C, with 45–65% relative humidity and a 12-hour light/dark cycle. They were utilized for parasite propagation, drug selection, parasite cloning.

### Plasmid construction and parasite transfection

The CRISPR/Cas9 plasmid pYCm was employed for genomic modification of the parasite^49^. To construct plasmid vectors for gene editing, 5′ and 3′ genomic sequences (300 - 600 bp) of the target genes were amplified as homology arms using specific primers (Table S2) and inserted into the designated restriction sites of pYCm. Pairs of oligonucleotides encoding single-guide RNAs (sgRNAs) (Table S2) were mixed, denatured at 95 °C for 3 min, annealed at room temperature for 5 min, and subsequently ligated into pYCm. The sgRNAs were designed to target the coding region of each gene (Table S2) using the PlasmoDB database. DNA fragments encoding HA tags were inserted in-frame between the left and right homology arms, fused to the gene of interest. For each gene, at least two sgRNAs were designed to target sites proximal to the C-terminal region. Infected red blood cells (iRBCs) were electroporated with 5 μg of plasmid using a Nucleofector 2b Device (Lonza, Germany). Transfected parasites were immediately injected intravenously into a naïve mouse and subjected to pyrimethamine treatment (7.0 mg/mL) beginning 24 h post-transfection. Under drug pressure, parasites carrying the transfected plasmids typically emerged within 5-7 days. For sequential modification, modified parasites were subjected to negative selection to eliminate the pYCm plasmid. 5-Fluorouracil (5-FC, Sigma-Aldrich, F6627) was dissolved in water (2.0 mg/mL) and administered to mice via drinking water in a light-protected bottle.

Naive mice receiving parasites with residual plasmids after pyrimethamine selection were subjected to 5-FC pressure until PCR confirmed plasmid elimination. All the genetically modified parasite lines generated in this study were listed in Table S3.

### Gametocyte induction in mouse

ICR mice were treated with phenylhydrazine (80 μg/g body weight; Sangon Biotech, China, cat# A600705-0025) via intraperitoneal injection. Three days after injection, the mice were infected with 6.0 × 10⁶ parasites by intravenous injection. Gametocytemia usually reached its peak on day 3 post-infection. Male and female gametocytes were quantified on Giemsa-stained blood smears. Sex-specific gametocytemia was calculated as the proportion of male or female gametocytes relative to parasitized erythrocytes.

### Gametocyte purification

Gametocytes were purified using the method described previously^50^with minor modifications. Briefly, mice were treated with phenylhydrazine 3 days before infection. From day 3 post-infection, sulfadiazine (20 mg/L; Acmec Biochemical, China, cat#S31660) was administered in drinking water to eliminate the asexual blood stage parasites. After two days of treatment with sulfadiazine, approximately 1 mL of gametocyte-containing mouse blood was collected from the orbital sinus and suspended in 6 mL gametocyte maintenance buffer (GMB, 137 mM NaCl, 4 mM KCl, 1 mM CaCl2, 20 mM glucose, 20 mM HEPES, 4 mM NaHCO3, 0.1% BSA, and pH 7.2). The 7 mL parasite sample was layered on top of a 2 mL 48% Nycodenz solution (27.6% w/v Nycodenz in 5 mM Tris-HCl, 3 mM KCl, 0.3 mM EDTA, and pH 7.2) in a 15 mL Falcon tube. After centrifugation at 1900 g for 20 min, the gametocytes were collected from the interface layer and washed twice with GMB for further use.

### Male gametocyte exflagellation assay *in vitro*

A 2.5 μL aliquot of mouse tail blood containing 4-10% gametocytemia was added to 100 μL exflagellation medium (RPMI 1640, 100 µM xanthurenic acid [XA; Tokyo Chemical Industry, Japan, cat#X0054], 2 U/mL heparin, pH 7.4). After 10 min of incubation at 22 °C, the exflagellation centers (ECs) and RBCs were counted in a hemocytometer under a light microscope. The percentage of RBCs containing male gametocytes was calculated from Giemsa-stained smears, and the exflagellation rate was calculated as ECs per 100 male gametocytes.

### Male gametes purification

Purification of male gametes was performed following the protocol for *P. berghei* male gamete isolation^33^. Purified gametocytes were activated in exflagellation medium for 15 min. The parasites were then centrifuged at 550 × g for 5 min, and this step was repeated twice.

Extracellular male and female gametes in the supernatant were collected by centrifugation at 10,000 × g for 7 min. The pelleted parasites were incubated at room temperature for 10 min to allow male gametes to swim into the supernatant. Subsequently, the supernatant was harvested, centrifuged at 12,000 × g for 10 min at 4 °C, and washed twice with PBS. The purity and quantity of the sample was determined by microscopy following Giemsa (Sigma-Aldrich, cat#, 48900-500ML-F) staining.

### Antibodies and antiserum

Primary antibodies include: rabbit anti-HA (Cell Signaling Technology, cat#3724S, 1:1000 for immunofluorescence IF), mouse antibody against both α-Tubulin I and α-Tubulin II (Sigma-Aldrich, cat#T6199, 1:1000 for IF). Secondary antibodies include: Alexa 555 goat anti-rabbit IgG (ThermoFisher Scientific, cat#A21428, 1:1000 for IF), Alexa 488 goat anti-mouse IgG (ThermoFisher Scientific, cat#A11001, 1:1000 for IF).

### Immunofluorescence assay

Purified parasites were fixed with 4% paraformaldehyde and transferred onto coverslips coated with poly-L-lysine (Sigma-Aldrich, cat# A-003-M) in a 24-well plate. Fixed cells were permeabilized with 0.1% Triton X-100 in PBS for 10 min, blocked with 5% BSA in PBS for 1 h at room temperature, and then incubated with primary antibodies diluted in 3% BSA-PBS for 1 h at room temperature. After three washes with PBS, the samples were incubated with fluorescently labeled secondary antibodies in 3% BSA-PBS for 1 h at room temperature. Hoechst 33342 was applied at a 1:5000 dilution in PBS for 15 min at room temperature.

Finally, the coverslips were washed, mounted in 90% glycerol, and sealed with nail varnish. All images were acquired using a Zeiss LSM 980 confocal microscope.

### Protein extraction and immunoblot

Parasites were lysed in RIPA buffer (Solaribio, cat#R0010) containing a protease inhibitor cocktail (MedChemExpress, cat#HY-K0010). After ultrasonication, the lysates were incubated on ice for 30 min and centrifuged at 14, 000 g for 10 min at 4℃. The supernatant was mixed with 6 volumes of pre-chilled (-20°C) pure acetone, precipitated for at least 30 min, and centrifuged at 14, 000 g for 30 min at 4°C. The supernatant was removed, and the protein pellet was stored at -80°C.

### Protein preparation and tryptic digestion

The extracted proteins were first solubilized in urea solution. Subsequently, dithiothreitol (DTT) was added to a final concentration of 10 mM, and the samples were thoroughly mixed and incubated at 37 °C for 60 min to reduce disulfide bonds. After reduction, iodoacetamide (IAA) was added to a final concentration of 20 mM, followed by incubation at 37 °C in the dark for 30 min to alkylate the free sulfhydryl groups. Excess IAA was quenched by adding DTT to a final concentration of 10 mM. The alkylated samples were then diluted with 50 mM ammonium bicarbonate buffer until the final urea concentration was below 2 M to ensure efficient tryptic digestion. Trypsin was added at an enzyme-to-protein ratio of 1:25 (w/w), and digestion was performed at 37 °C for 16 h. The enzymatic reaction was terminated by adding formic acid to a final concentration of 5%. The digested peptide samples were subsequently dried using vacuum centrifugation at 37 °C for approximately 2 h.

### Peptide desalting and LC-MS/MS analysis

The dried peptide mixtures were resuspended in 200 μL of loading buffer containing 0.5% trifluoroacetic acid (TFA) and 5% acetonitrile (ACN) in ultrapure water. Peptides were purified using Pierce C18 spin columns (Thermo Fisher Scientific) according to the manufacturer’s protocol. After desalting, the peptide samples were dried and reconstituted in 10 μL of 0.1% formic acid (FA) for subsequent LC-MS/MS analysis. LC-MS/MS analysis was performed on an Orbitrap Eclipse Tribrid mass spectrometer (Thermo Fisher Scientific) coupled with an EASY-nLC 1200 nano-flow liquid chromatography system. Peptide separation was achieved using a self-packed C18 analytical column (75 μm inner diameter × 20 cm length) packed with ReproSil-Pur 120 C18-AQ particles (1.9 μm, 120 Å). The peptides were separated at a flow rate of 300 nL/min using water containing 0.1% FA as mobile phase A and 80% ACN containing 0.1% FA as mobile phase B. A 120 min linear gradient was applied as follows: 5–10% B over 10 min, 10–35% B over 95 min, 35–40% B over 12 min, followed by an increase to 90% B within 3 min and maintained for 7 min. Mass spectrometric acquisition was conducted using a data-dependent acquisition (DDA) strategy. Full MS scans were acquired over an m/z range of 350–1500 at a resolution of 60,000, followed by MS/MS scans with a resolution of 15,000. Precursor ions were isolated using a 1.6 m/z isolation window and fragmented by higher-energy collisional dissociation (HCD) with a normalized collision energy of 30%. The cycle time was set to 2 s, and MS/MS acquisition was initiated from an m/z value of 120.

### Database searching and protein identification

The raw LC-MS/MS files were analyzed using Proteome Discoverer software (version 2.4.1.15, Thermo Scientific) with a standard proteomic database searching workflow. MS/MS spectra were searched against the UniProt^51^ reference proteome database of *Plasmodium yoelii*. The database search was performed with a minimum peptide length of 6 amino acids and a maximum of two missed cleavages allowed. Peptide-spectrum matches (PSMs) and peptide identifications were filtered using a false discovery rate (FDR) threshold of <1%. The precursor ion mass tolerance was set to 10 ppm, while the fragment ion mass tolerance was set to 0.02 Da.

### Transmission electron microscopy

Purified parasites were fixed with 4% glutaraldehyde in 0.1 M phosphate buffer at 4 °C overnight and subsequently washed three times with PBS. The samples were then post-fixed with 1% osmium tetroxide for 2 h and washed three times with PBS. Dehydration was carried out using a graded acetone series. After embedding and sectioning, thin sections were stained with uranyl acetate and lead citrate. Images of the parasites were acquired using a Hitachi HT-7800 electron microscope.

### Cryo-ET and Cryo-EM sample preparation

The R 2/1 EM grids (R2/1, Au, 200 mesh, Quantifoil) were glow-discharged for 80 s using the PELCO easiGlow Glow Discharge Cleaning System (Ted Pella) before sample preparations. *Plasmodium* male gametes were loaded onto the glow discharged EM grids and vitrified using the Vitrobot Mark IV (Thermo Fisher Scientific). For cryo-EM sample, both sides were blotted for 5 s with a force of 4; for cryo-ET sample, only one side was blotted for 5 s with a force of 8. All grids were stored in liquid nitrogen until data collection.

### cryo-FIB milling

Briefly, the grids were transferred into a Helios 5 Cryo-FIB/SEM (Thermo Fisher Scientific). To reduce the charging effect, a protective layer of organometallic platinum was deposited using the gas injection system (GIS), followed by sputter coating. Rough milling was performed with a current of 0.5 nA in the region of interest to produce an initial lamella approximately 1.5 μm thick. The current was then progressively reduced, and a final polishing step was carried out at 24 pA, resulting in a lamella with a thickness of 150–200 nm. The lamellae were then stored in liquid nitrogen for subsequent data collection.

### Cryo-EM and cryo-ET data collection

Cryo-EM data was collected on a 300 kV Titan Krios transmission electron microscope (Thermo Fisher Scientific) equipped with a Falcon 4 camera and a Gatan energy filter. A total of 32,738 movies were collected over five sessions at a nominal magnification of 105,000× (calibrated pixel size: 1.21 Å), with a defocus range of −0.5 to −2.0 μm. Each micrograph received a total dose of 61 e^-^/Å^2^, with exposure time of ∼10 s.

Cryo-ET data of *Plasmodium* male gametes were acquired on a Titan Krios (Thermo Fisher Scientific, G4) equipped with a Gatan energy filter and a Falcon 4 direct electron detector, using Tomography 5 software for automated dose-symmetric acquisition. For samples at 15 mpa, images were collected with a dose-symmetric tilt scheme from −60° to 60° at a 2° step, at a nominal magnification of 105,000× (calibrated pixel size: 1.21 Å) and a defocus range of −1.5 to −4.0 μm; a total of 265 tilt series were acquired. For samples at 2 mpa and 8 mpa, the tilt scheme ranged from +62° to −38° with a 2° increment (beginning at a pretilt of +12°), at a nominal magnification of 53,000× (calibrated pixel size: 2.37 Å) and a defocus range of −3.0 to −5.0 μm; a total of 404 tilt series were collected. Each tilt series in both datasets received a total dose of approximately 110 e⁻/Å².

### Cryo-EM image processing of the DMT

Image preprocessing was performed in CryoSPARC^52^. Motion correction and contrast transfer function (CTF) estimation were performed on all 32,738 movies. To curate a high-quality dataset, micrographs with predicted resolutions worse than 10 Å were discarded.

Particle selection for the 9+2 axonemal doublet microtubules (DMTs) was performed using GisSPA^34^, a template-matching based software. To generate template for GisSPA, ∼950 k DMT particles were manually picked at an 8 nm interval. These particles were used to resolve a 48 nm repeat DMT template map at 5.08 Å resolution after 3D classification and local refinement (described below). After getting the DMT particles using GisSPA, most of the false positives were removed by 3D classification with cylindrical masks on B tube and inner junction (IJ) in RELION3^53^. To compensate the missing particles, Hermite interpolation was applied, yielding 1,540,960 particles at an 84 Å interval. Finally, all particles at 1x binning were used to generate a 3.96 Å resolution DMT map to preclude the possibility of reference bias.

The selected particles were then imported to CryoSPARC and extracted with a box size of 152 pixels at 4x binning. A local refinement was performed with a rotation search extent of 15° and a shift search extent of 21 Å to refine particle poses. To resolve the 48-nm repeat DMT structure, we employed a two-step 3D classification at 2x binning, with alignment disabled and tubulin signal subtracted to increase particle periodicities. First, a cylindrical mask covering the outer dynein arm (ODA) was applied, yielding three well-resolved classes (each ∼33%) that revealed 24-nm repeat features. Subsequently, a cylindrical mask focused on the MIPs at the IJ was used, yielding two classes (each ∼50%) exhibiting the 48-nm periodicity. The 48-nm particles were shifted by ±48 nm to compensate missing particles, and duplicates were removed. Finally, a local refinement produced the consensus 48-nm DMT map at 4.64 Å resolution.

To improve the structural details of 48-nm repeat map, local refinements with 30 overlapping cylindrical masks covering different regions were performed. All resulted in improved maps (with the best resolution of 3.28 Å), which were fitted back into consensus map to generate a final composite map. To improve the maps of ODA and ODA docking complex (DC), local refinements were performed in the 24-nm particles with the focused masks, yielding maps with resolution of 4.29 Å (ODA-DC) and 7.03 Å (ODA).

### Tomogram reconstruction and visualization

Tilt series were motion-corrected using MotionCor2^54^ and then reconstructed using IMOD^55^ or AreTomo^56^. For visualization, high-quality tomograms were processed using IsoNet2^57^ for denoising and missing-wedge compensation. Three-dimensional renderings of the axonemes were generated based on the subtomogram averaged results obtained the following steps.

### Subtomogram averaging of the DMT and central pair

DMTs were manually traced in IMOD following their polarity. Subtomograms were picked at an 8-nm interval using the interpolation function in RELION2 and cropped at 6x binning via WARP^58^. Initial particle poses were determined through 3D refinement in RELION2, with constraints applied to the tilt and psi Euler angles and a shift search range limited to 6 pixels (∼4.3 nm). Subsequent local refinement and M^59^ refinement at 2x binning yielded an 8-nm repeat DMT map at 6.3 Å resolution.

To resolve the 48-nm repeat DMT map, we applied a two-step 3D classification strategy which is analogous to the cryo-EM processing described above. The resulting 48-nm particles were refined to a resolution of 7.9 Å. To resolve the 96-nm repeat DMT map, a cylindrical mask covering the MAP densities near A02-A03 protofilament was used to perform 3D classification at 4x binning, yielding two well-revolved classes (each ∼50%). Subsequently, a 3D classification with local refinement focusing on radial spoke (RS) was applied, and one class containing intact RS was selected and refined to generate the 96-nm repeat DMT map. Central pairs were manually traced in IMOD following their polarity. Subtomograms were extracted at an 8-nm interval using the interpolation function in RELION2 and cropped at 12x binning via WARP. Initial particle poses were determined through 3D refinement in RELION2, with the tilt and psi Euler angles constrained and a shift search range limited to 6 pixels (∼8.6 nm). The result displayed a 16 -nm repeat structure. Duplicate particles were removed, and rot Euler angles were unified using scripts.

As the relative position of C1 and C2 microtubules exhibit flexibility, they were resolved separately using masks specific to either C1 or C2. For C1, a round of 3D classification with a cylindrical mask covering the left-side MAPs was performed, which yielded a 32-nm repeat. In contrast, the C2 exhibits 16-nm periodicity despite several 3D classification tasks were tried. Finally, particle centers were shifted to the centers of C1 and C2 respectively, followed by independent 3D refinements to improve the final resolution.

### Model building and protein identification

Atomic models for 48-nm repeat DMT were de novo built using CryoAtom^43^ automatically. The results were then manually refined in Coot^60^.

For the identification of microtubule inner proteins unique to the *Plasmodium*, the protein sequences of *Plasmodium yoelii* 17X from UniPort (Proteomes ID: UP000072874) were applied for CryoAtom. The candidate proteins were manually evaluated through our LC-MS/MS results and AlphaFold2^61^ predicted models.

### Ortholog search using protein BLAST

To search the orthologs of IDA, RS1 components, NME7, CFAP161 and RIB72, the amino acid sequences of those proteins in *Chlamydomonas reinhardtii* were used as the query sequences. The BLAST searches were performed in NCBI^62^, UniProt and PlasmoDB^63^ databases. For NCBI searches, PSI-BLAST (Position-Specific Iterated BLAST) was selected with the organism restricted to *Plasmodium yoelii* (taxid:5861). For UniProt and PlasmoDB searches, the organism was restricted to *P.yoelii* 17X, while all other parameters were kept at default settings. The top 10 hits from each search were manually evaluated based on their predicted structures (AlphaFold2) and transcriptomic profiles from the Malaria Cell Atlas^64^. Proteins lacking similar structures or not mainly expressed in male gametocyte were excluded.

## Acknowledgments

We are grateful to the Cryo-EM Platform of Peking University and Changping Laboratory for data collection, and the National Centre for Protein Sciences at Peking University for technical assistance. This work is funded by, the National Natural Science Foundation of China (32600007, 22474005), the Beijing Natural Science Foundation (JQ24031 to Q.G.), Ministry of Science and Technology of the People’s Republic of China (2024YFC2309700, 2024YFA1306200 and 2025YFA1309300). Q.G. is supported by the SLS-Qidong Innovation Fund. The authors declare no competing financial interests.

## Data availability

The representative tomogram of gametocyte at 2, 8, 15 minutes post activation has been deposited in the Electron Microscopy Data Bank (EMDB) under the accession code EMD-83137, EMD-83138, EMD-83139. The subtomogram averaging maps of 96-nm repeating DMT, C1 and C2 complex of central pair have been deposited in the EMDB under the accession codes EMD-83153, EMD-83154, EMD-83155. Cryo-EM coordinates and density maps have been deposited in the Protein Data Bank (PDB) and EMDB under the accession codes PDB 44XD, EMD-83079.

## Notes

### Competing Interest Statement

The authors have declared no competing interest.

