## Supplemental File for "Structural streamlining of axonemal doublet microtubules in *Plasmodium* male gametes"

### Supplementary

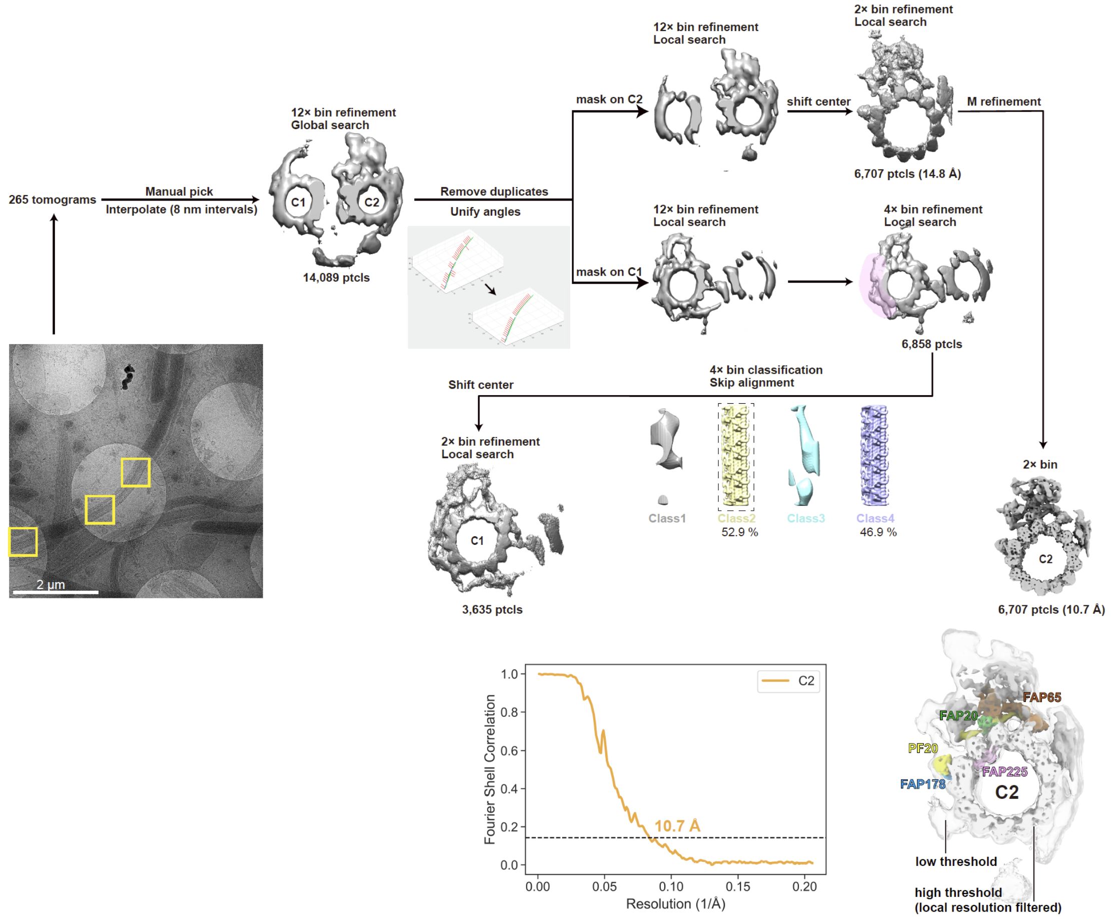


**Fig. S1 Subtomogram averaging processing pipeline of the *Plasmodium* central pair.**

Schematic overview from particle picking to final map generation. Fourier shell correlation (FSC) curves for the 16-nm repeat C2 tubule refinement are displayed. The proteins homologous to *C.reinhardtii* in C2 are annotated.

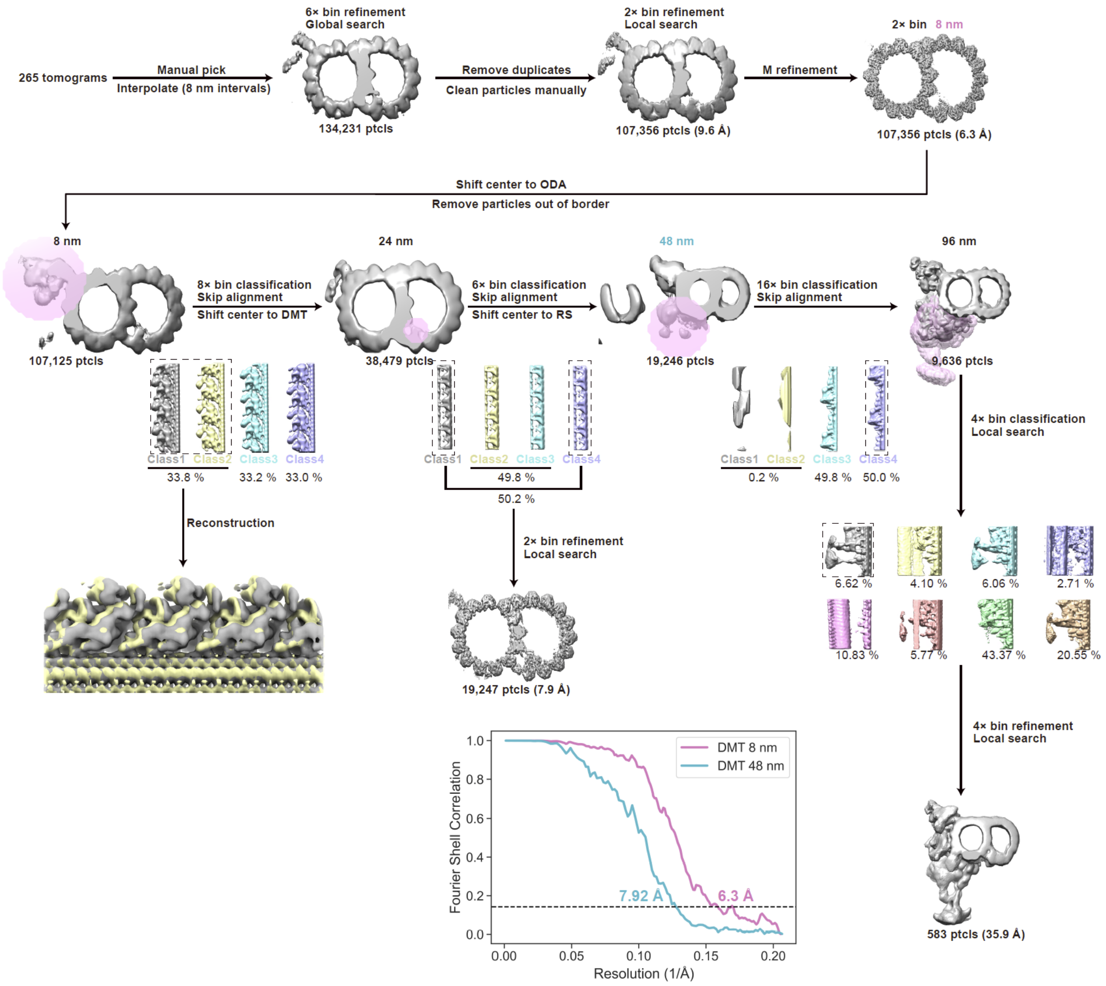


**Fig. S2 Subtomogram averaging processing pipeline of the *Plasmodium* DMT.** Schematic overview from particle picking to final map generation. Fourier shell correlation (FSC) curves for the 48-nm and 96-nm repeat refinement are displayed. (RS: radial spoke, ptcls: particles)

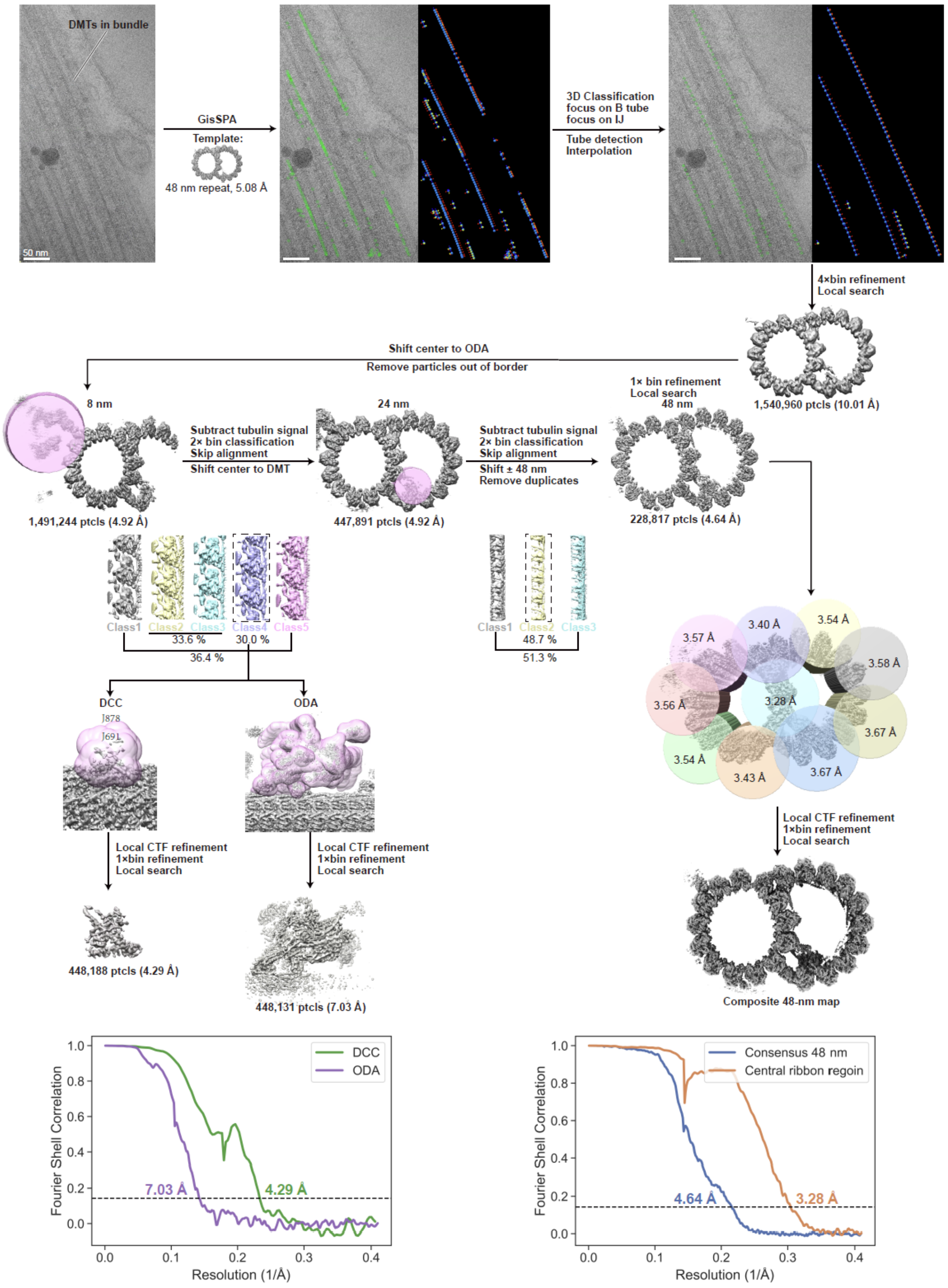


**Fig. S3 Cryo-EM processing pipeline of the *Plasmodium* DMT.** Schematic overview from particle picking to composite map generation. Fourier shell correlation (FSC) curves for the dock complex calmodulin (DCC), outer dynein arm (ODA), central ribbon focused refinement and consensus refinement are displayed. (IJ: inner junction, ptcls: particles)

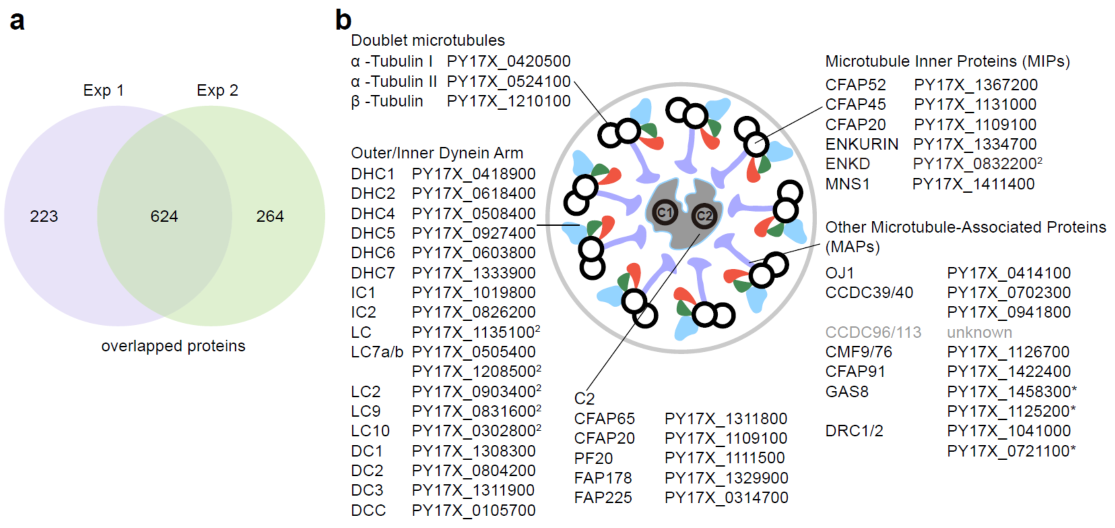


**Fig. S4 Protein identification based on mass spectrometry and cryo-ET/EM density maps**

**a**. Venn diagrams of proteins detected by mass spectrum (MS) in *Plasmodium* male gametes from two independent experiments.

**b**. Proteins detected via analysis of MS and experimental density map. Proteins are grouped by region. Protein names (left) and corresponding gene IDs (right) are listed. Superscript numbers indicate proteins detected only in the corresponding experimental batch. Asterisks indicate low sequence homology.

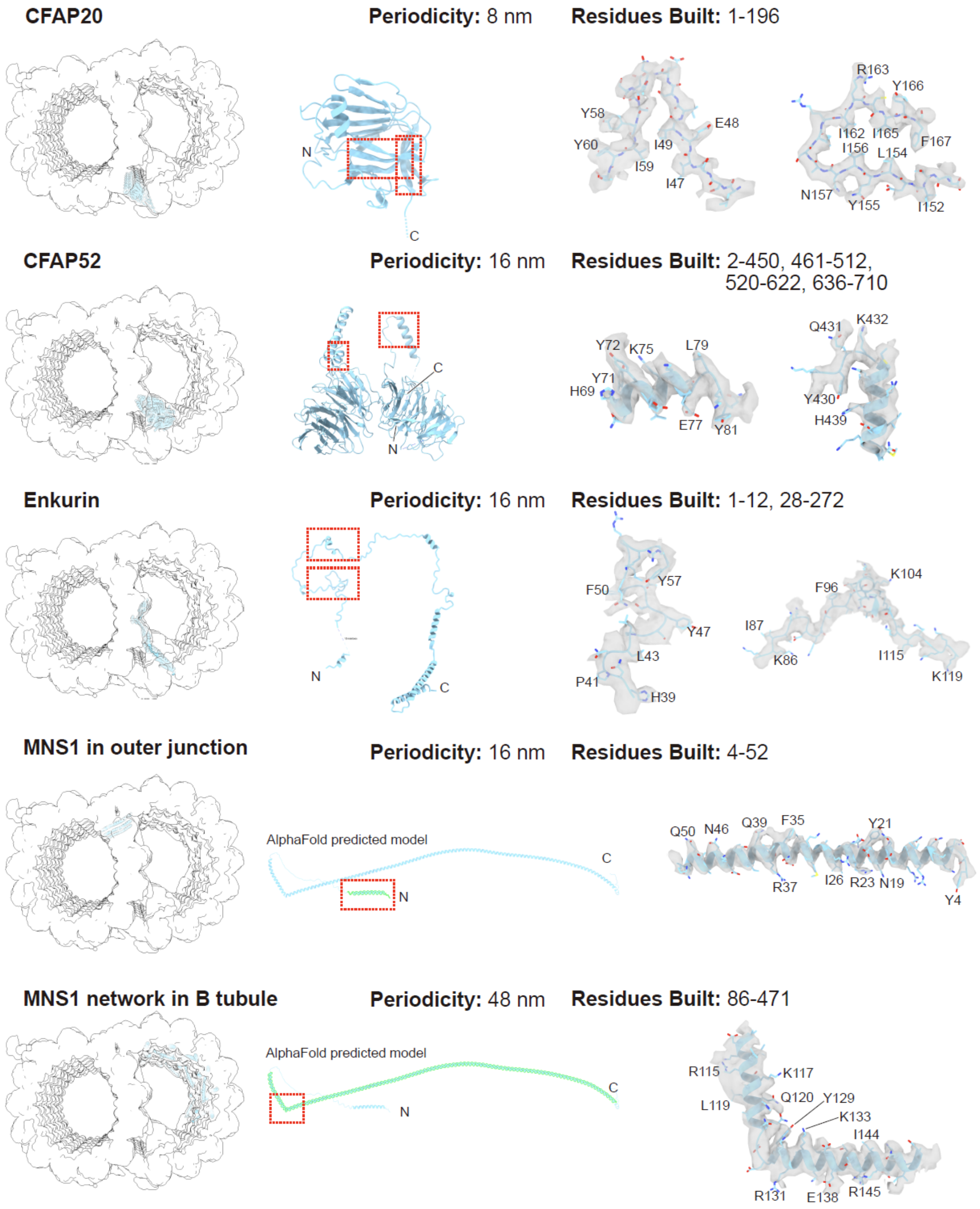
CFAP20
Periodicity: 8 nm
Residues Built: 1-196
N
C
Y58
Y60
I59
I49
I47
E48
R163
Y166
I162
I165
I156
L154
F167
N157
Y155
I152
CFAP52
Periodicity: 16 nm
Residues Built: 2-450, 461-512, 520-622, 636-710
C
N
Y72
K75
L79
Y71
H69
E77
Y81
Q431
K432
Y430
H439
Enkurin
Periodicity: 16 nm
Residues Built: 1-12, 28-272
N
C
F50
Y57
Y47
L43
P41
H39
F96
K104
I87
K86
I115
K119
MNS1 in outer junction
Periodicity: 16 nm
Residues Built: 4-52
AlphaFold predicted model
C
N
Q50
N46
Q39
F35
Y21
R37
I26
R23
N19
Y4
MNS1 network in B tubule
Periodicity: 48 nm
Residues Built: 86-471
AlphaFold predicted model
C
N
R115
K117
L119
Q120
Y129
K133
I144
R131
E138
R145

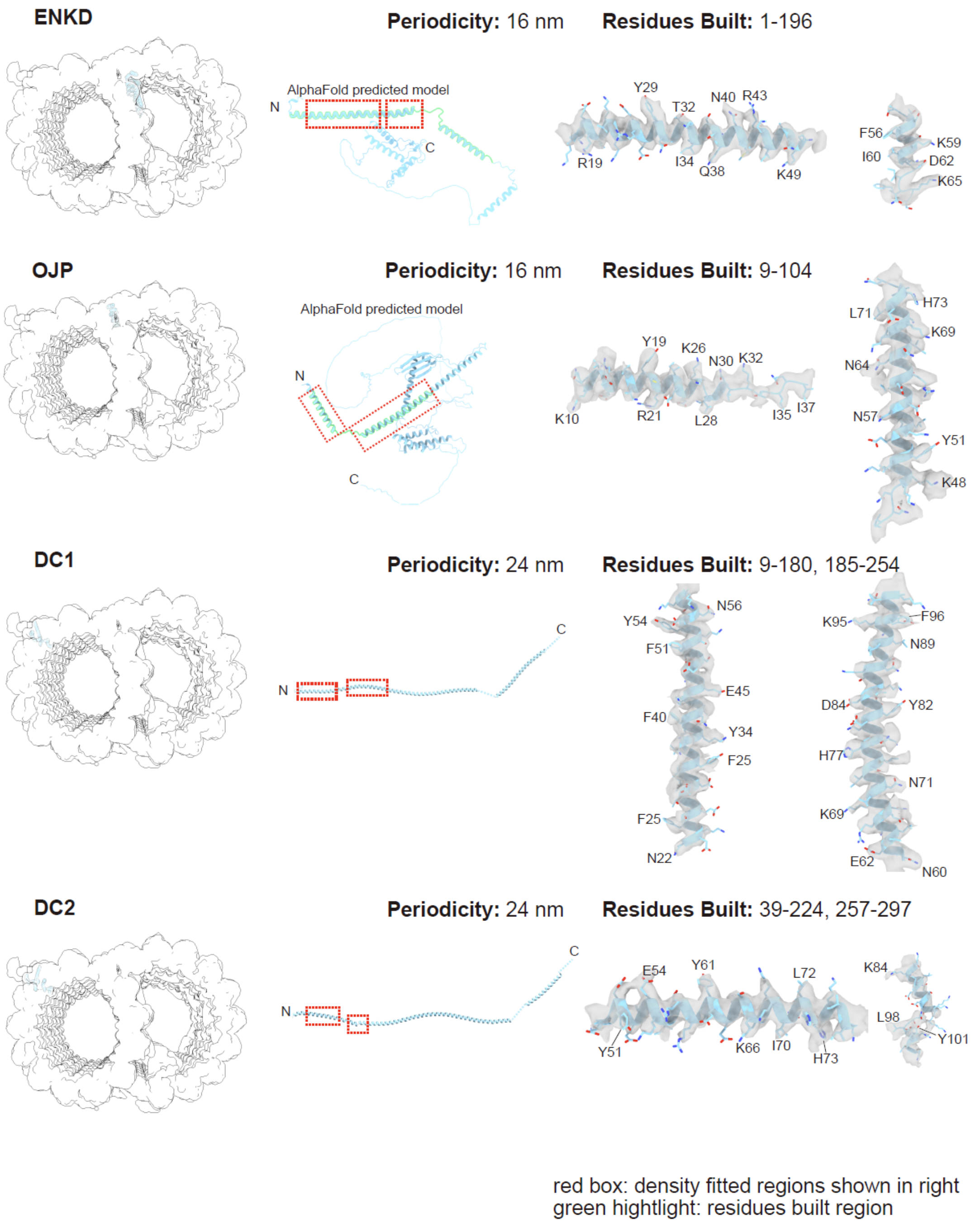


**Fig. S5 Location, atomic models and representative density of microtubule inner proteins.** Atomic models include either AlphaFold-predicted models or models built into the experimental density map. Regions corresponding to the experimental density map are highlighted in green. The representative density maps shown are indicated by the red boxes.

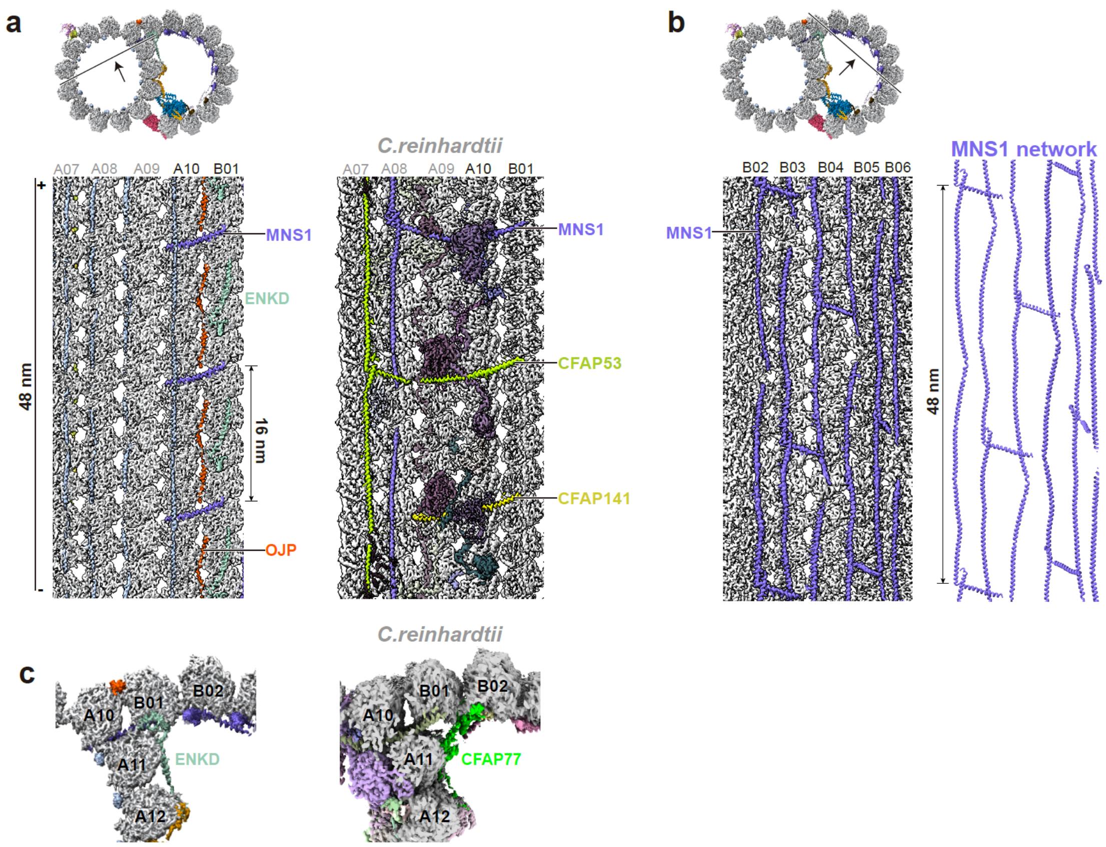


**Fig. S6 Architecture of the MNS1 in DMTs.**

**a**. Longitudinal views of the MNS1 on protofilaments of A10 and B01. MNS1, CFAP53 and CFAP141 are spaced by 16 nm in *Chlamydomonas*, whereas the sites of CFAP53 and CFAP141 are occupied by MNS1 in *Plasmodium.*

**b**. Longitudinal views of the MNS1 in B tubules. The MNS1 forms a network architecture with periodicity of 48 nm.

**c**. Cross-sectional views of the outer junction in *Plasmodium* (left), with ENKD, and in *Chlamydomonas* (right), with CFAP77.

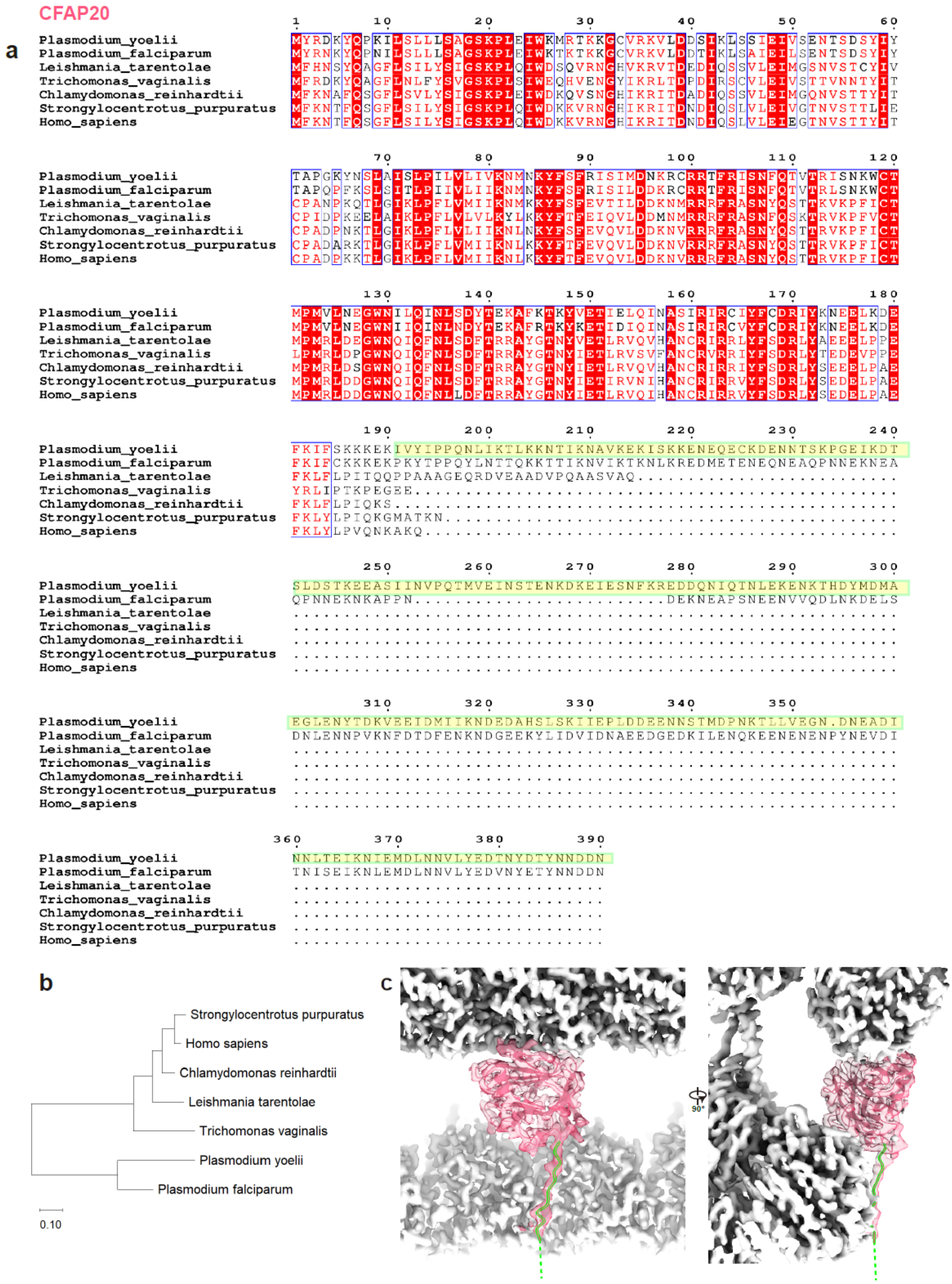


**Fig. S7 Comparative sequence and structural analyses of CFAP20**

**a**. Multiple protein sequence alignment of CFAP20 showing a *Plasmodium*-specific sequence extension (highlighted in yellow and green). The alignment includes orthologs from six other species and was generated using ClustalW with Gonnet protein weight matrix.

**b**. Phylogenetic tree of CFAP20. Evolutionary analysis of CFAP20 via the Maximum Likelihood method.

**c**. Atomic model of CFAP20 (red) bound to tubulins (white). *Plasmodium*-specific extension was highlighted in yellow and green, corresponding to the sequence highlighted in (a).

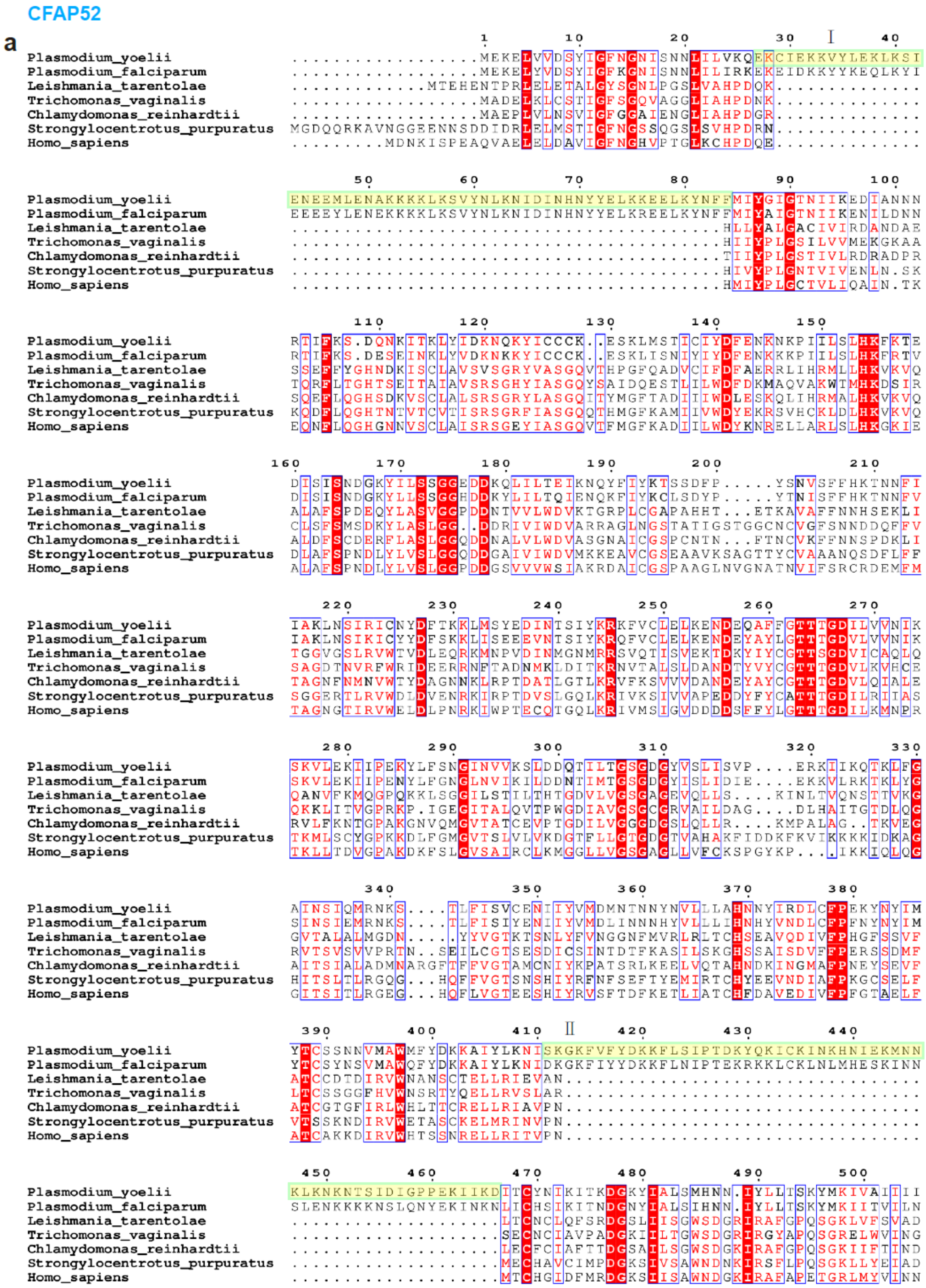
CFAP52
a
Plasmodium_yoelii
Plasmodium_falciparum
Leishmania_tarentolae
Trichomonas_vaginalis
Chlamydomonas_reinhardtii
Strongylocentrotus_purpuratus
Homo_sapiens
I
II

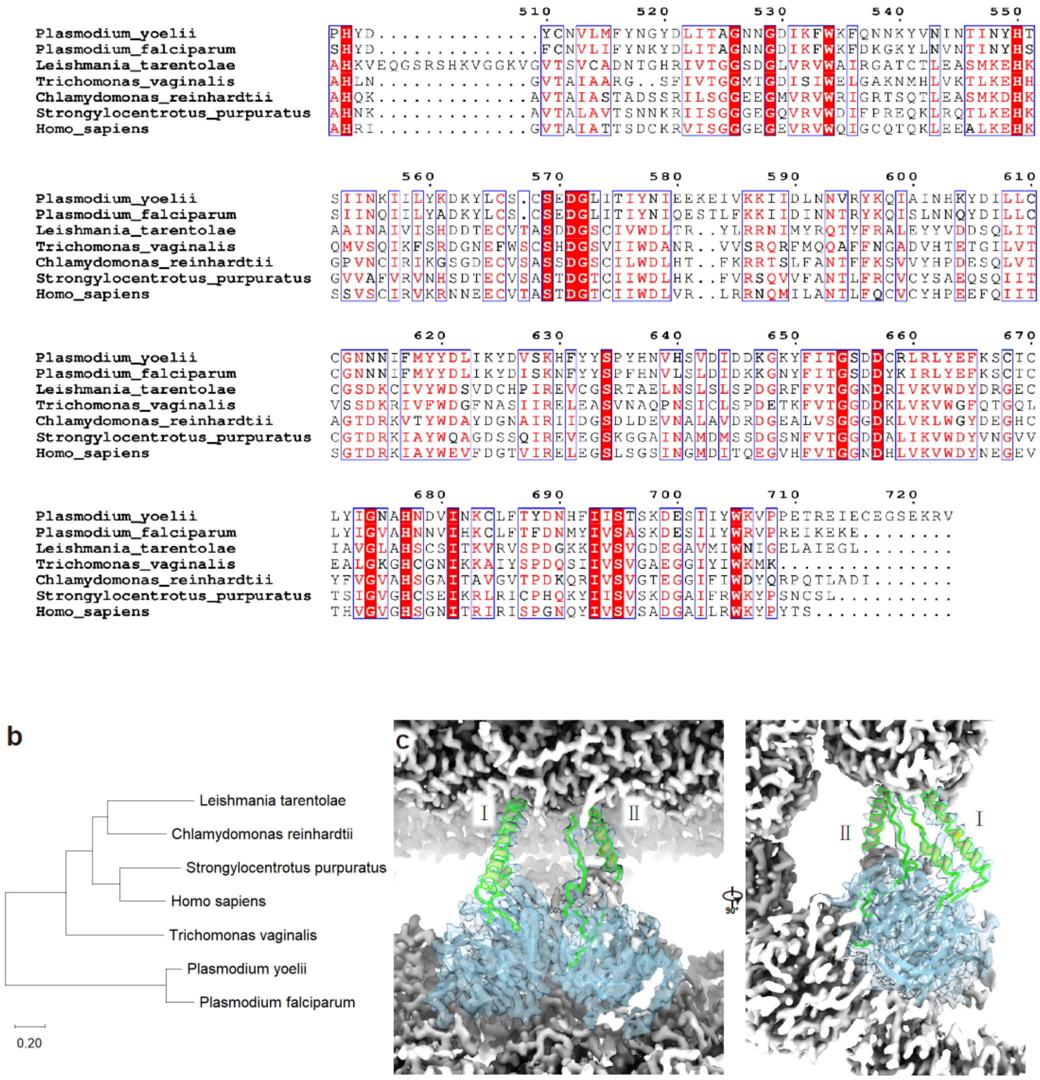


**Fig. S8 Comparative sequence and structural analyses of CFAP52**

**a**. Multiple protein sequence alignment of CFAP52 showing a *Plasmodium*-specific sequence extension (highlighted in yellow and green). The alignment includes orthologs from six other species and was generated using ClustalW with Gonnet protein weight matrix.

**b**. Phylogenetic tree of CFAP52. Evolutionary analysis of CFAP52 via the Maximum Likelihood method.

**c**. Atomic model of CFAP52 (blue) bound to tubulins (white). *Plasmodium*-specific extension was highlighted in yellow and green, corresponding to the sequence highlighted in (a).

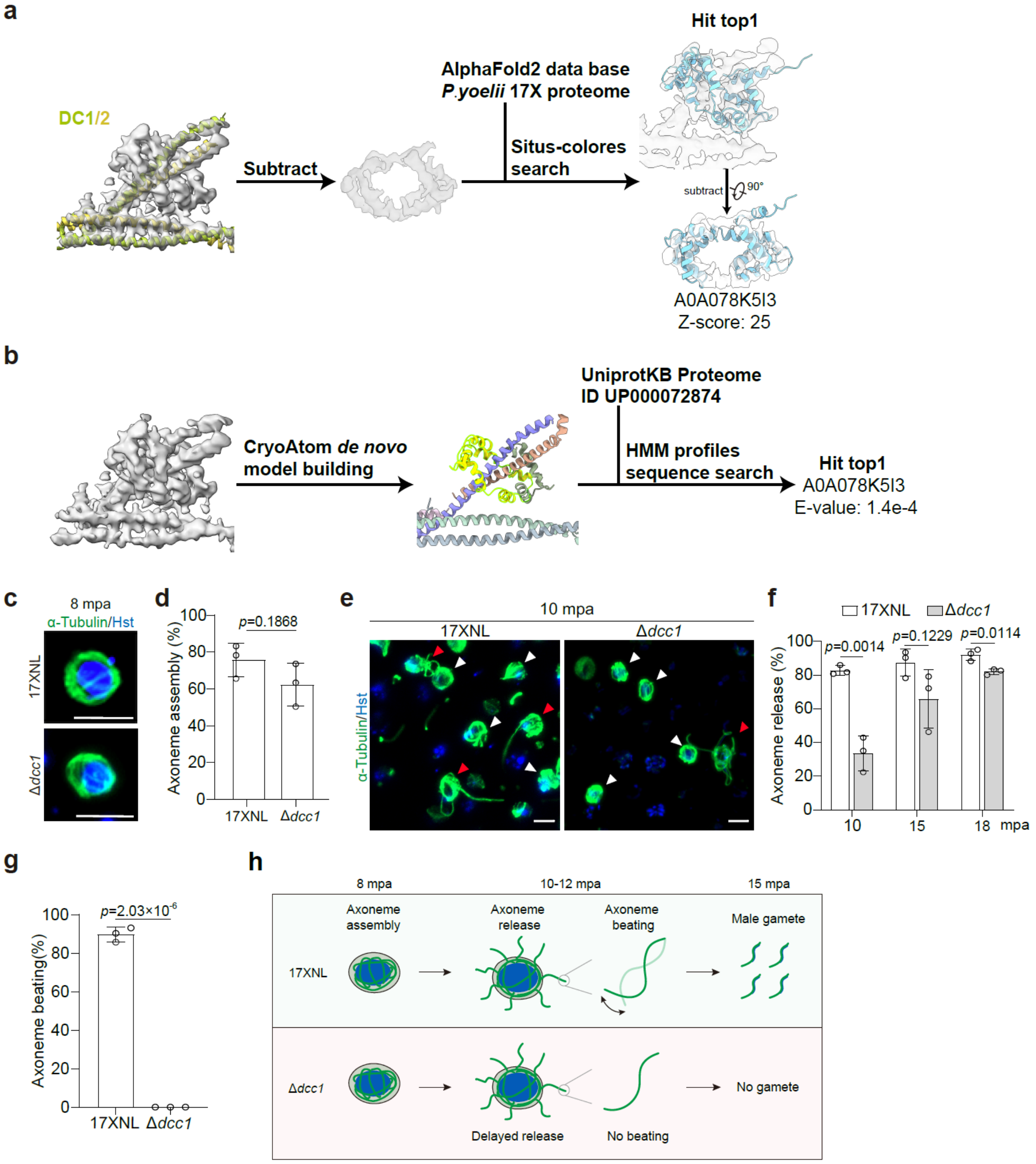


**Fig. S9 Identification of calcium-binding protein in ODA docking complex.**

**a**. The densities of DC1/2 were subtracted, leaving the density of calcium-binding protein for model fitting using Situs software[1]. Candidate models were obtained from AlphaFold2 database of *P.yoelii* 17 X and fitted into the density map.

**b**. CryoAtom software[2] was used for *de novo* model building based on the density map. The resulting atomic model of calcium-binding protein (colored in yellow) was used to perform sequence search using the hidden Markov model (HMM) profiles. Protein sequences used for the HMM search were downloaded from the UniProt[3] database.

**c**. IFA showing the formation of cytoplasmic axonemes in gametocytes at 8 mpa. Parasites were stained with the α-Tubulin antibodies and Hoechst 33342. Scale bars, 5 μm.

**d**. Quantification of male gametocytes with axoneme assembly in C. Mean ± SD from three independent experiments; two-tailed t-test.

**e**. IFA showing axoneme release in gametocytes at 10 mpa. Gametocytes with unreleased (white) and releasing (red) axonemes are indicated by arrows.

**f**. Quantification of axoneme release in the fully assembled male gametocytes at 10, 15, and 18 mpa. Mean ± SD from three independent experiments, two-tailed t-test.

**g**. Quantification of male gametocytes with actively beating axonemes at 10 mpa by bright-field microscopy. Mean ± SD from three independent experiments; two-tailed t-test.

**h**. Cartoon illustrating the defects in axoneme release, axoneme beating, and male gamete formation upon DCC deficiency during male gametogenesis.

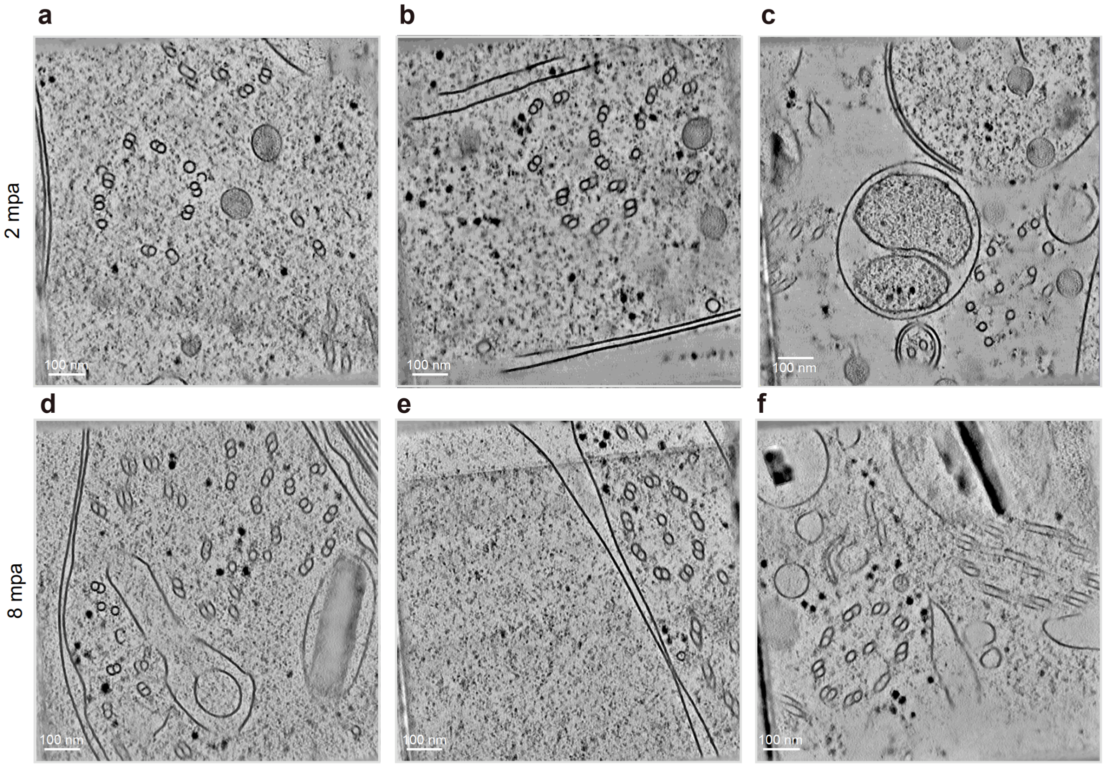


**Fig. S10 Cryo-ET of *P. yoelii* male gametocytes during gametogenesis**

**a-f**. Tomograms showing *P. yoelii* male gametocytes activated for 2 minutes (a-c) and 8 minutes (d-f), respectively.

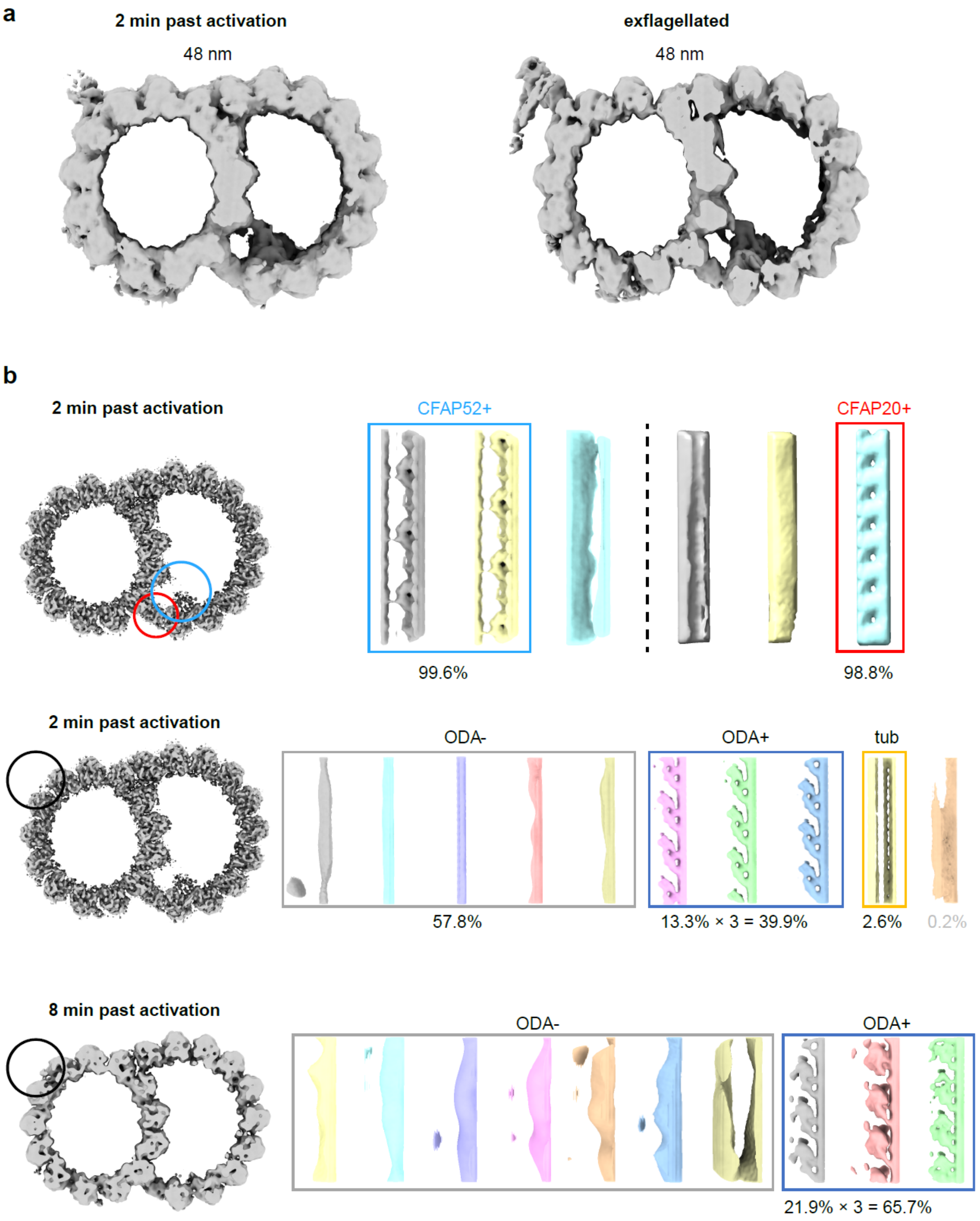


**Fig. S11 Development of microtubule doublets during male gametogenesis**

**a**. Subtomogram-averaged structures of DMTs from male gametocytes two minutes after activation (left) and exflagellated male gametes (right).

**b**. Three-dimensional classification reveals CFAP20, CFAP52 and ODA assembly at different time points. CFAP52+, CFAP20+, ODA+, classes containing corresponding proteins; tub, classes in which tubulin occupies the ODA-binding sites; mix, classes containing ODA and other heterogeneous particles.

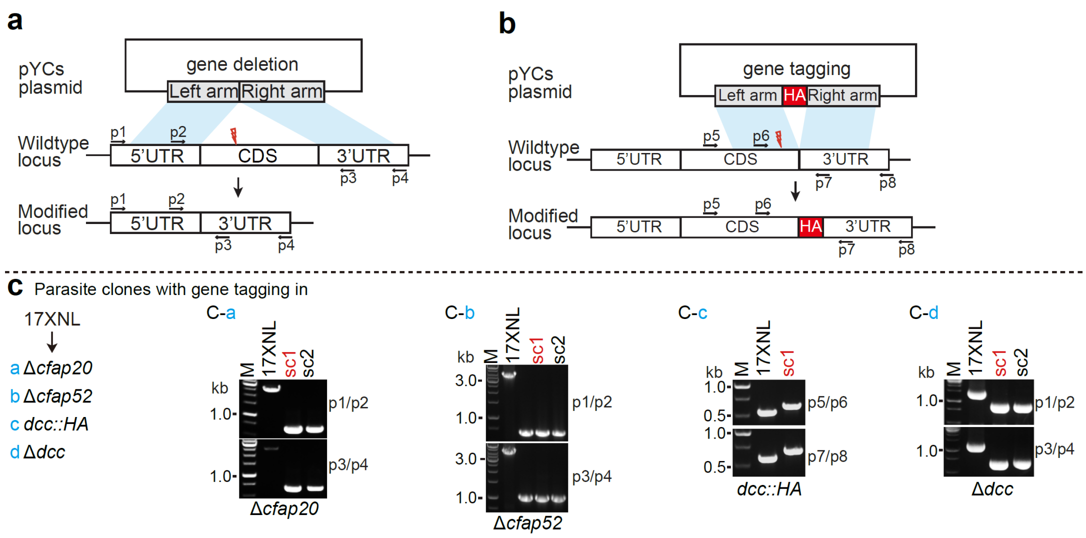


**Fig. S12 Genotyping of genetically modified parasites in this study**

**a-b**. Schematic representation for CRISPR-Cas9-mediated gene modification, including gene tagging (a) and gene deletion (b) via double-cross homologous recombination.

**c**. For each modification, both 5' and 3' homologous recombination events were detected by the genotyping PCR (see the primers in the Table S2) to confirm successful integration of the homologous templates. Two independent experiments yielded similar results. Usually, 1-2 parasite single clones (sc) were obtained after limiting dilution, and one clone (indicated with red letter) was used for further phenotype and gene functional analysis.

| | #1 composite map of *Plasmodium yoelii* DMT, 48 nm repeat (EMDB-83079) (PDB 44XD) | #2 Subtomogram averaging of *Plasmodium yoelii* DMT, 96 nm repeat (EMDB-83153) | #3 Subtomogram averaging of *Plasmodium yoelii* central pair apparatus C1, 32 nm repeat (EMDB-83154) | #4 Subtomogram averaging of *Plasmodium yoelii* central pair apparatus C2, 16 nm repeat (EMDB-83155) |
|---|---|---|---|---|
| **Data collection and processing** | | | | |
| Magnification | 105,000× | 105,000× | 105,000× | 105,000× |
| Voltage (kV) | 300 | 300 | 300 | 300 |
| Electron exposure (e–/Å$^2$) | 61 | 110 | 110 | 110 |
| Defocus range (μm) | -0.5 to -2.0 | -2.5 to -4.0 | -2.5 to -4.0 | -2.5 to -4.0 |
| Pixel size (Å) | 1.21 | 4.84 | 2.42 | 2.42 |
| Symmetry imposed | C1 | C1 | C1 | C1 |
| Initial particle images (no.) | 1,540,960 | 134,231 | 14,105 | 14,105 |
| Final particle images (no.) | 228,817 | 583 | 3,634 | 6,707 |
| Map resolution (Å) | 3.54 (30 maps ave.) | 35.90 | 12.1 | 10.7 |
| FSC threshold | 0.143 | 0.143 | 0.143 | 0.143 |
| Map resolution range (Å) | 3.28 to 3.76 | None | None | None |
| **Refinement** | | | | |
| Initial model used (PDB code) | None | | | |
| Model resolution (Å) | 2.3 | | | |
| FSC threshold | 0.5 | | | |
| Model resolution range (Å) | None | | | |
| Map sharpening *B* factor (Å$^2$) | 27.2 to 46.3 | | | |
| Model composition | | | | |
| Non-hydrogen atoms | 1,065,152 | | | |
| Protein residues | 136,175 | | | |
| Ligands | 420 | | | |
| *B* factors (Å$^2$) | | | | |
| Protein | 122.55 | | | |
| Ligand | 103.84 | | | |
| R.m.s. deviations | | | | |
| Bond lengths (Å) | 0.003 | | | |
| Bond angles (°) | 0.657 | | | |

| | |
|---|---|
| Validation | |
| MolProbity score | 1.69 |
| Clashscore | 10.60 |
| Poor rotamers (%) | 0.17 |
| Ramachandran plot | |
| Favored (%) | 97.19 |
| Allowed (%) | 2.78 |
| Disallowed (%) | 0.03 |

**Table S1.** Cryo-EM data collection, refinement and validation statistic.

| **Primers for gene knockout** | | | | | |
|---|---|---|---|---|---|
| **Gene name** | **Gene ID** | **Modified strain** | **Gene size /deleted gene size (bp)** | **Left homologous template** | |
| | | | | **Forward primer** | **Reverse primer** |
| *Δcfap20* | PY17X_1109100 | 17XNL | 2227/2227 | CGGGGTACCCACTTTATACGTATACTAA | CATGCCATGGATTTAAAGACGTACTTAA |
| *Δcfap52* | PY17X_1367200 | 17XNL | 3304/3346 | CCCAAGCTTGCATGTGTAATGTAAAAAATAAT | CATGCCATGGGGTTATATAGATCTTTCCTTTCC |
| *Δdcc* | PY17X_0105700 | 17XNL | 474/474 | CCCAAGCTTGGTAGGAAAGCAAACGTAG | CATGCCATGGTTTTAGCTAGCTATTTTTTTATAT |
| **Gene name** | **Gene ID** | **Right homologous template** | | **Target site of sgRNA** | |
| | | **Forward primer** | **Reverse primer** | **Oligo (Forward)** | **Oligo (Reverse)** |
| *Δcfap20* | PY17X_1109100 | CCGCTCGAGGGGGTCCATATAAGATGGAA | CCCCTTAAGACATGACCTTATCACAGATT | TATTGGAAAATGAGAACAAAAAAA | AAACTTTTTTTGTTCTCATTTTCC |
| *Δcfap52* | PY17X_1367200 | CCGCTCGAGGAAAATCACAAAAAAATATGCAGA | CCCCTTAAGCGAGGAGTATCAAATACCAG | TATTGTTCGAACAATGTTATGGCA | AAACTGCCATAACATTGTTCGAAC |
| *Δdcc* | PY17X_0105700 | CCGCTCGAGAGATTATATATCTCATAATAATTTA | CCCCTTAAGTAAATGGGGACCACAATTAA | TATTGAAAATAATAGAGATTGTAA | AAACTTACAATCTCTATTATTTTC |
| **Primers for PCR-genotyping parasite with gene knockout** | | | | | |
| **Gene name** | **Gene ID** | **P1** | **P2** | **P3** | **P4** |
| *Δcfap20* | PY17X_1109100 | ATGCACTAGTCATATATGTT | TGTTGCTTGGCTTGTTATGT | ATGACATACTAAGCACACAC | ATAAATGGGTTTAACCTTGT |
| *Δcfap52* | PY17X_1367200 | GCACACCGTTTATTTTTCTA | GACCTGAAATATGGCTCACA | TTAAGCCCAATTTAAGCTCC | GCCCAAATTATGTAAATCGA |
| *Δdcc* | PY17X_0105700 | GAGAATATACGTCTGCACAT | TAAATGGGGACCACAATTAA | CTTTACTCACACATATAAGAC | ATAAGAACATAAATTTGTATCAC |
| **Primers for gene tagging** | | | | | |
| **Gene name** | **Gene ID** | **Modified strain** | **Tag (name and location)** | **Left homologous template** | |
| | | | | **Forward primer** | **Rerverse primer** |
| *dcc::HA* | PY17X_0105700 | 17XNL | C-terminal HA | CCCAAGCTTATGAAGAATTATGTGATGATC | CATGCCATGGAATATTCCATATAGCATTTTTA |
| **Gene name** | **Gene ID** | **Right homologous template** | | **Target site of sgRNA** | |
| | | **Forward primer** | **Reverse primer** | **Oligo (Forward)** | **Oligo (Reverse)** |
| *dcc::HA* | PY17X_0105700 | CCGCTCGAGAGATTATATATCTCATAATAATTTA | CCCCTTAAGTAAATGGGGACCACAATTAA | TATTGTATATACATATTCCGTGTGC | AAACGCACACGGAATATGTATATAC |
| **Primer sequence for PCR-genotyping parasite with gene tagging** | | | | | |
| **Gene name** | **Tag (name and location)** | **Modified strain** | **Gene ID** | **P1** | **P2** |
| *dcc::HA* | C-terminal HA | 17XNL | PY17X_0105700 | CTCTGATGAAATAAACAATTTAGAG | AAATCGGTAATTCCATTTTGA |
| **Gene name** | **Tag (name and location)** | **Modified strain** | **Gene ID** | **P3** | **P4** |
| *dcc::HA* | C-terminal HA | 17XNL | PY17X_0105700 | GACAACGAATTTAAGGAGG | AAAAGTGACCATAATATTCATC |

**Table S2.** Primers and oligonucleotides used in this study.

| **Strain** | **Description** | | | **Resource** |
|---|---|---|---|---|
| 17XNL | *Plasmodium yoelii* 17XNL strain | | | NIH |
| **Parasite with gene deletion** | **Gene ID** | **Parental strain** | **Description** | **Resource** |
| Δ*cfap20* | PY17X_1109100 | 17XNL | Deleted the whole coding sequence of *cfap20* in the 17XNL parasite | This study |
| Δ*cfap52* | PY17X_1367200 | 17XNL | Deleted the whole coding sequence of *cfap52* in the 17XNL parasite | This study |
| *dcc::HA* | PY17X_0105700 | 17XNL | C-terminally tagged with HA of *dcc* in the 17XNL parasite | This study |
| Δ*dcc* | PY17X_0105700 | 17XNL | Deleted the whole coding sequence of *dcc* in the 17XNL parasite | This study |

**Table S3.** List of genetically modified parasite strains used in this study.
